# Geometric cues stabilise long-axis polarisation of PAR protein patterns in *C. elegans*

**DOI:** 10.1101/451880

**Authors:** Raphaela Geßele, Jacob Halatek, Laeschkir Würthner, Erwin Frey

## Abstract

In the *Caenorhabditis elegans* zygote, PAR protein patterns, driven by mutual anatagonism, determine the anterior-posterior axis and facilitate the redistribution of proteins for the first cell division. Yet, the factors that determine the selection of the polarity axis remain unclear. We present a reaction-diffusion model in realistic cell geometry, based on biomolecular reactions and accounting for the coupling between membrane and cytosolic dynamics. We find that the kinetics of the phosphorylation-dephosphorylation cycle of PARs and the diffusive protein fluxes from the cytosol towards the membrane are crucial for the robust selection of the anterior-posterior axis for polarisation. The local ratio of membrane surface to cytosolic volume is the main geometric cue that initiates pattern formation, while the choice of the long-axis for polarisation is largely determined by the length of the aPAR-pPAR interface, and mediated by processes that minimise the diffusive fluxes of PAR proteins between cytosol and membrane.

## Introduction

Cell polarisation is a crucial process in development^1^. Well studied examples include localisation of bud sites in *Saccharomyces cerevisiae*^2^, apico-basal asymmetry in mammalian epithelial cells^3^, and the asymmetric placement of the first cell division in the *Caenorhabditis elegans* zygote^4^. A key question in such systems is how the correct polarity axis is established and robustly maintained.

In *C. elegans*, the anterior-posterior axis of the embryo is determined in the fertilised egg by a polarised distribution of PAR (partitioning defective) proteins^4–6^. Immediately before the establishment of polarisation begins, the future anterior PARs (aPARs) cover the cell cortex uniformly, while posterior PARs (pPARs) are cytoplasmic^7^. After fertilisation, the sperm-donated centrosome induces contraction of the actomyosin network, which leads to cortical flows that displace cortical aPARs anteriorly, allowing cytoplasmic pPARs to bind in the posterior zone^8–11^; see Fig. 1*A*. Once these two PAR domains have formed (during the ‘establishment phase’) and have thereby established the anterior-posterior axis, they persist for several minutes through the ‘maintenance’ phase until cell division^5, 7^.

**Figure 1:**
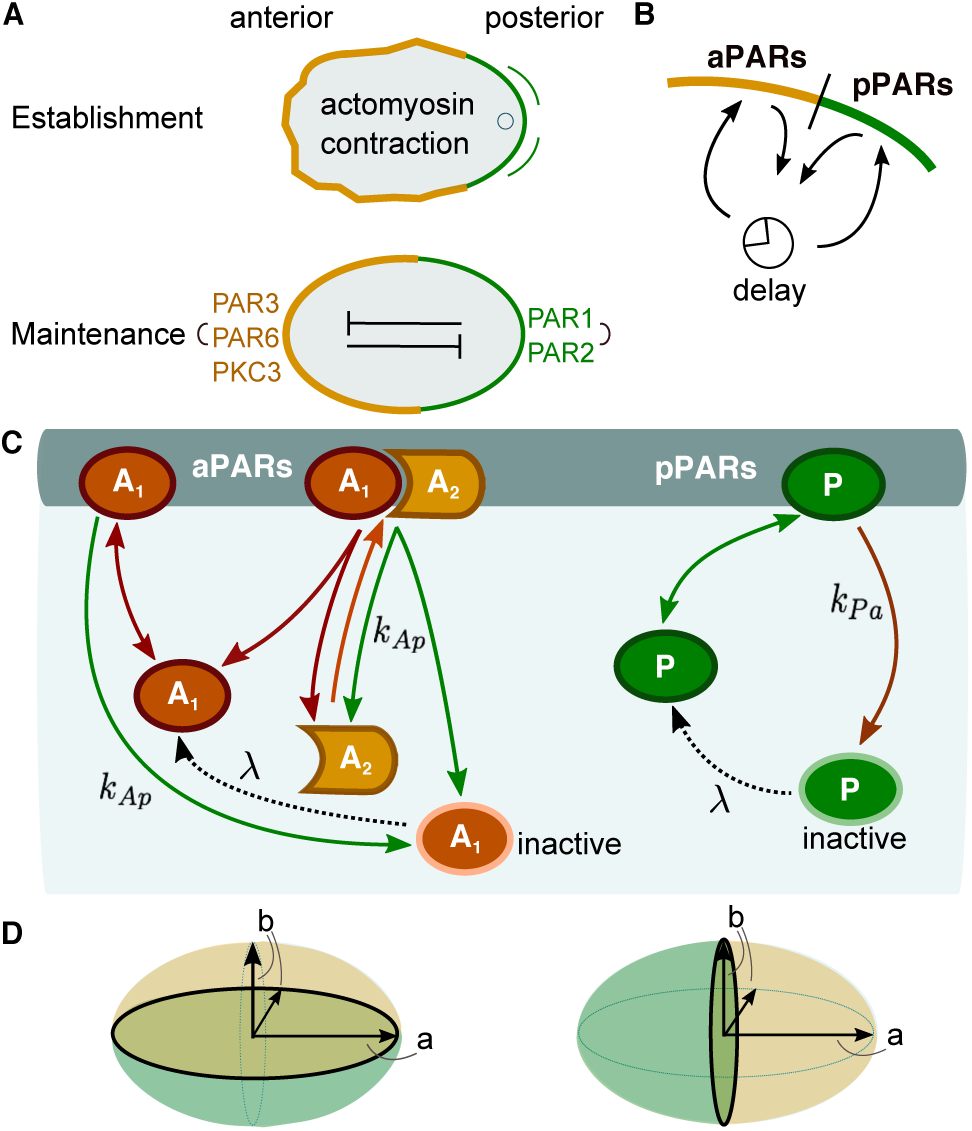
Biological background and model network. (*A*) Cell polarisation in the *C. elegans* embryo during the establishment (top) and maintenance (bottom) phases; sketch adapted from Ref.^5^. (*B*) Illustration of protein flux between cytosol and membrane. As proteins detach from the membrane when phosphorylated, they cannot immediately rebind to the membrane. There is therefore an intrinsic delay before dephosphorylation permits rebinding. (*C*) The biochemical reaction network is comprised of two mutually antagonistic sets of proteins, aPARs and pPARs. Dephosphorylated (active) *A*_1_ and *P* attach to the membrane with rates 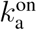 and 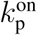, respectively. Both active proteins may also detach spontaneously from the membrane with rates 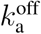 and 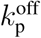, respectively. *A*_1_ acts as a scaffold protein: Once bound to the membrane it recruits *A*_2_ with rate *k*_d_ and forms a membrane-bound hetero-dimeric aPAR complex *A*_12_. The hetero-dimer *A*_12_ may itself spontaneously detach from the membrane with rate 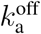 and dissociate into *A*_2_ and active *A*_1_. Membrane-bound *A*_1_ and *A*_12_ can also be phosphorylated by *P* with rate *k*_Ap_[*P*], thereby initiating dissociation of the aPAR complex and release of aPAR proteins into the cytosol. While reattachment of the scaffold protein *A*_1_ is delayed by the requirement for dephosphorylation (reactivation), detached *A*_2_ can be recruited to the membrane by membrane-bound *A*_1_ immediately. Similarly, *P* is phosphorylated by the hetero-dimer *A*_12_ at rate *k*_Pa_[*A*_12_], and is consequently released as inactive *P* into the cytosol. In the same way as *A*_1_, also *P* must be dephosphorylated before it can bind again to the membrane. For simplicity, we take identical dephosphorylation (reactivation) rates λ for inactive *A*_1_ and *P*. The ensuing reaction-diffusion equations are provided in the Method section and a table listing the values of the rate constants can be found in 1. (*D*) Sketch of the cell’s geometry: Prolate spheroid with long axis *a* and short axis *b*, and with short- (left) and long-axis (right) polarisation.

Several independent in vivo experiments on *C. elegans* have demonstrated that maintenance of PAR protein polarity is independent of an intact actomyosin network^7, 11–15^. Rather, it appears that the entry of the sperm and the following contractions of the cortical actomyosin serve as a temporal trigger for the rapid establishment of the PAR protein pattern^9, 13, 16^. However, experimental observations also suggest that while the rapid establishment and perfect position of anterior-posterior PAR domains are the result of an interplay between mechanical, hydrodynamical and biochemical mechanisms, polarisation is nevertheless robustly established (albeit with some delay) when various mechanical and hydrodynamical mechanisms are eliminated.^10, 11, 17–19^. To disentangle and understand these distinct mechanisms one needs to investigate the mechanism of self-organised polarisation by the biochemical PAR protein network. Based on the fact that aPAR and pPAR proteins mutually drive each other off the membrane by phosphorylation^20^, and that this antagonism promotes formation of distinct domains on the membrane^10, 21, 22^, previous studies have outlined how self-organisation of PAR proteins maintain polarisation until cell division^**?**, 15, 16^. These studies showed that basic features of PAR protein polarisation can be explained by minimal reaction-diffusion models. However, as these models used a simplified one-dimensional geometry and assumed that cytosolic proteins are homogeneously distributed, the effect of cell geometry was disregarded and the distinction between long and short axis was lost. Thus, how the long axis is selected for polarisation and subsequently maintained, and in a broader context, which features of a reaction-diffusion system are responsible for axis selection remain open questions.

To answer these questions we draw on previous studies of other intracellular pattern-forming protein systems which revealed that even the typically rather fast cytosolic diffusion does not eliminate protein gradients in the cytosol^23–26^. As a consequence, protein patterns are generically sensitive to cell geometry through coupling between processes in the cytosol and on the membrane. In particular, it was predicted^23, 24^ that delayed reattachment to the cell membrane (e.g., due to cytosolic nucleotide exchange) is key to geometry sensing. Indeed, recent experimental studies support the idea that axis selection depends on the interplay between reaction kinetics and cellular geometry^25^.

These results suggest that the protein dynamics in the cytoplasm of the *C. elegans* embryo may also influence the selection of the long over the short axis during polarity maintenance. In order to investigate axis alignment, we developed a reaction-diffusion model of the PAR protein dynamics. As in previous studies^9, 15, 27^, a central element in our model is mutual displacement of membrane-bound aPARs and pPARs by phosphorylation. However, in contrast to earlier models^9, 28^, we do not use effective nonlinearities but strictly biomolecular reactions based on mass-action law kinetics, e.g. by explicitly modelling the formation of PAR protein complexes. Importantly, we also account for the delay caused by the need for reactivation of detached PAR proteins by cytosolic dephosphorylation, thus introducing the generic feature of a biochemical activation-deactivation cycle.

Our extended reaction-diffusion model in realistic cell geometry reveals that the dynamics of the phosphorylation-dephosphorylation cycle of PAR proteins is crucial for long-axis polarisation. Without this additional feature, the biochemical network of PAR proteins would not lead to robust polarisation along the long axis but instead exhibit a strong tendency to first polarise along the short axis, and polarisation would not re-align within a time that corresponds to a typical time before cell division. Furthermore, the extended model enables us to characterise the roles of mutual antagonism (phosphorylation) and overall protein numbers in robust long-axis polarisation: while the phosphorylation rates determine how distinctively one polarisation axis is selected over the other, relative protein numbers primarily affect the robustness of pattern formation as a whole.

Most importantly, our analysis indicates that these findings can be generalised beyond the specific model for the PAR system: axis selection is based on the generic dependence of intracellular pattern-forming processes on the local ratio of membrane surface to cytosolic volume and on the cell geometry via the length of the interface between the two different protein domains. Broadly speaking, the membrane-to-bulk ratio determines the likelihood that a given protein will reattach to the membrane quickly after detachment into the cytosol and the interface length affects both the establishment and maintenance of long-axis polarisation.

## Results

### Model

The aPAR set of proteins comprises PAR-3, PAR-6, and the atypical protein kinase PKC-3. Only complexes containing PKC-3 can phosphorylate pPARs, thereby disabling their membrane--binding capacity^21, 29^. How trimeric complexes consisting of PAR-3, PAR-6 and PKC-3 actually form is not fully understood. The evidence so far suggests that PAR-6 acts as a linker between PKC-3 and PAR-3, which can itself bind directly to the membrane^30–33^. In the absence of PAR-6, PKC-3 freely diffuses in the cytosol^34, 35^. In the reaction network upon which our mathematical model is based, we simplify the formation of trimeric complexes to the formation of a complex consisting of two effective species of aPARs: *A*_1_ and *A*_2_ (Fig. 1*C*). The first species, *A*_1_, models the membrane binding function of PAR-3, thus we also refer to it as a scaffold protein. The second species, *A*_2_, corresponds to a complex of PAR-6 and PKC-3. It is assumed to be recruited by scaffold proteins *A*_1_ that are already bound to the membrane, thereby forming hetero-dimers *A*_12_ on the membrane. These complexes can then phosphorylate membrane-bound pPARs, which initiates their release into the cytosol in a phosphorylated (inactive) state.

As with aPARs, there are different pPAR species, PAR-1 and PAR-2. While it is known that PAR-2 binds directly to the membrane, and PAR-1 phosphorylates PAR-3, it remains unclear whether PAR-2 also helps to maintain anterior-posterior polarity by excluding aPAR complexes from the membrane^7, 20^. However, PAR-2 is required for posterior binding of PAR-1^36^ and PAR-2 exclusion from the membrane by PKC-3 is essential for proper restriction of pPARs to the posterior^21^. In view of the remaining uncertainties we refrain from distinguishing between different species and effectively treat the pPARs as a single species *P* (Fig. 1*C*). *P* phosphorylates membrane-bound *A*_1_ and *A*_12_, which triggers their subsequent detachment as a phosphorylated (inactive) species into the cytosol.

Our model also accounts for protein dephosphorylation reactions in the cytosol. This creates deactivation-reactivation cycles, as proteins that were phosphorylated (deactivated) on the membrane are thereby reactivated for membrane binding (Fig. 1*B, C*). For simplicity, the reactivation (dephosphorylation) rate λ is assumed to be identical for cytosolic pPARs (*P*) and aPARs (only *A*_1_). The ensuing reaction-diffusion equations are given in the Method section Equations (7-18).

We approximate the natural shape of a *C. elegans* embryo by a prolate spheroid with semi--axis lengths *a* = 27*µm* and *b* = 15*µm* (see Fig. 1 *D*) ^9^. Here, *a* is the distance from centre to pole through a focus along the symmetry axis, also called the semi-major axis, while *b* is the equatorial radius of the spheroid, which is called the semi-minor axis. The boundary and interior of the ellipse represent the cell membrane and cytosolic volume, respectively.

### Dephosphorylation plays a key role for axis determination

For mutually antagonistic protein interactions, protein domains are separated by an interface at which mutually induced membrane detachment dominates^9, 15, 16^. For its maintenance proteins that have detached from the membrane must be replaced, otherwise the antagonistic interaction between the proteins would deplete either aPARs or pPARs from the membrane. As the protein interactions are mass-conserving, maintenance requires that detached proteins quickly rebind, unless the cytosolic reservoir of proteins is large enough for them to be replenished directly. This suggests that an interface can best be maintained locally in those membrane regions where rebinding to the membrane after detachment is most likely.

The likelihood of rebinding depends on the availability of cytosolic proteins for binding, which depends on the interplay between the local cell geometry and the time required for reactivation of detached proteins by dephosphorylation (Fig. 2). The ratio of available membrane surface to cytosolic volume is highest at cell poles and lowest at mid-cell. How this local cell geometry affects protein rebinding depends on the dephosphorylation time: a longer reactivation time implies that a protein that detached in a phosphorylated state from the membrane will on average diffuse farther away from the membrane before it can be reactivated and reattaches. The corresponding reactivation length is estimated as

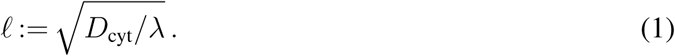

**Figure 2:**
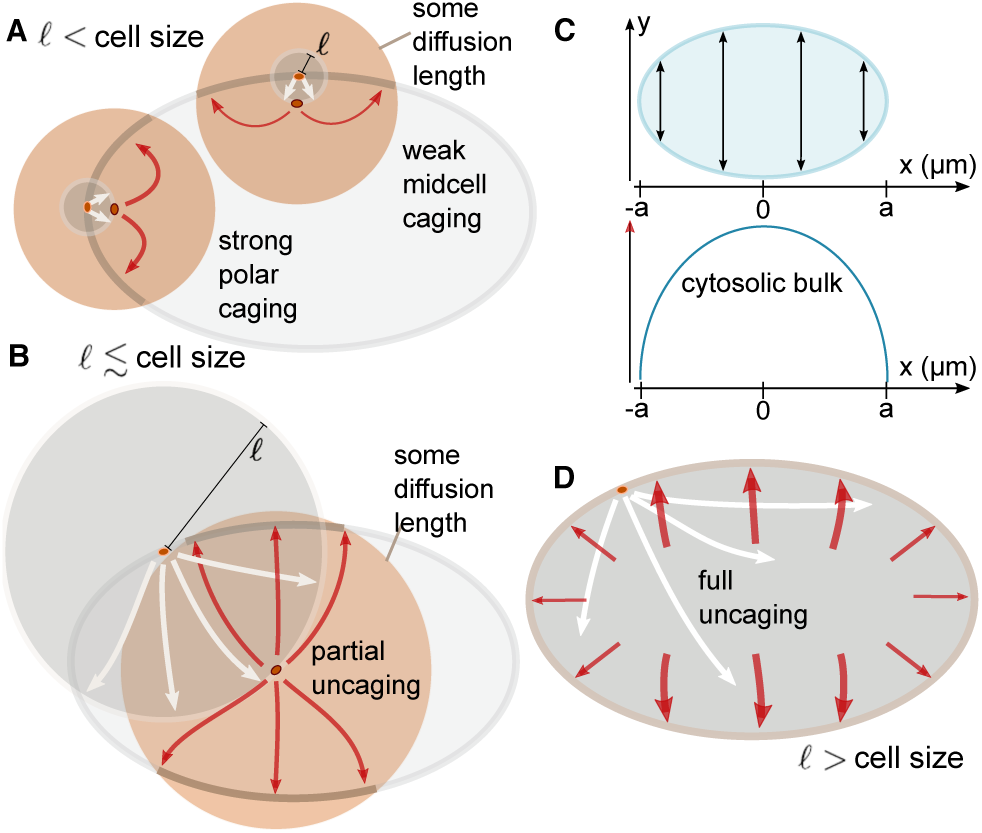
Role of dephosphorylation in axis determination. (*A, B*, and *D*): A protein is shown in the elliptical cell firstly at its phosphorylation and detachment site on the membrane and then at the point of its reactivation. The reactivation length gives an average radius (gray circles) how far from the detachment point a protein travels before reactivation. The orange circles around the reactivated protein and the associated arrows sketch some diffusion distance corresponding to a time interval Δ*t* following reactivation, i.e. during this time interval the protein can reattach to the membrane. (*A*) If the reactivation length *ℓ* (radius of gray circle) is small compared to the cell size, the local membrane surface to cytosolic volume ratio strongly affects the position at which detached proteins reattach. Due to the reactivation occurring close to the membrane, within some time interval Δ*t* following reactivation a protein that detaches from a cell pole is more likely to reattach near that same cell pole than a protein detaching from mid-cell is to reattach at mid-cell. Hence, dynamics that are based on membrane-cytosol cycling (such as antagonistic reactions that maintain an interface) are enhanced at the cell poles. (*B*) As the reactivation length *ℓ* approaches the length of the cell, this effect of geometry becomes weaker, and detaching proteins become increasingly unconstrained by the position of detachment (uncaged). (*C*) Illustration of the distribution of cytosolic bulk proteins along the long-axis. The elliptical cell and the cytosol height is depicted as a function of *x*, where the *x*-axis aligns with the long axis (top). The amount of cytosolic bulk proteins for each *x* varies from the poles to mid-cell as illustrated (bottom). (*D*) This effect of cell geometry is completely lost if the reactivation length *ℓ* exceeds the length of the cell. Hence, detached proteins become uniformly distributed throughout the cell before reactivation occurs. In that case, most will re-encounter the membrane near mid-cell after reactivation, since a delocalised protein will most likely be found in the mid-cell area.

To see how this diffusion length affects protein dynamics, consider a protein with a short inactive (phosphorylated) phase, such that *ℓ* is significantly smaller than the cell length *L* = 2*a* (Fig. 2*A*). Then, proteins are likely to be dephosphorylated fast and can therefore rebind very soon after phosphorylation-induced detachment. Since the local ratio of membrane surface to cytosolic volume at the cell poles is larger than at mid-cell, these proteins are more likely to reencounter the membrane in the polar zone which translates into higher polar reattachment (after reactivation), i.e. proteins remain caged at the cell poles (Fig. 2*A*). Conversely, proteins that detached from the membrane at mid-cell have more cytosolic volume available than those that detached at the poles and, thus, are less likely to re-encounter the membrane and rebind there (Fig. 2*A*). This heuristic picture suggests that for *ℓ* ≪ *L* domain interfaces preferentially form at the cell poles and hence cell polarity will be established along the short-axis. If dephosphorylation requires more time, *ℓ* increases and the effect of local membrane curvature is attenuated (Fig. 2*B*). Ultimately, when *ℓ* > *L*, proteins can be considered as uniformly distributed throughout the cytosol for the next attachment event (Fig. 2*D*). Therefore, reactivated proteins are more likely to attach at mid-cell, where the accumulated density along the long-axis (or, equivalently, the ratio of cytosolic volume to membrane area) is highest (Fig. 2*C*). This implies that an interface between different protein domains will establish itself at mid-cell and cells will become polarised along the long-axis for large enough reactivation length *ℓ*.

In summary, if cell polarisation is induced by antagonistic protein interaction, we expect long-axis polarisation to be favoured only if the delay resulting from the inactive phase is sufficiently long. Moreover, our analysis suggests that relative protein numbers affect axis selection, as the global availability of an abundant protein species attenuates the effect of cell geometry associated with the activation-deactivation cycle.

In the heuristic arguments outlined above, we tacitly considered a single position along the interface between the PAR domains. In general, however, the length of the interface may also play an important role in determining the orientation of the axis ultimately selected, as one expects energetic costs for interface establishment and maintenance to scale with its length. In the following we will analyse the system’s dynamics in a two-dimensional as well as in a three-dimensional cell geometry; an analysis of a simplified rectangular geometry would actually be misleading (Supplementary Note 3). Furthermore, the analysis in two and three dimensions enables us to disentangle the effects due to the membrane-to-bulk ratio and interface length in polarisation establishment and maintenance. Note that in a two-dimensional ellipse the interface between the domains reduces to a point, such that all geometric effects can be solely attributed to the membrane-to-bulk ratio.

### Growth rates of long versus short-axis polarisation

To put the above heuristic reasoning concerning the role of membrane-to-bulk ratio on a firm basis, we first performed a mathematical analysis in two-dimensional elliptical geometry, building on previous investigations of intracellular pattern formation ^23, 24^.

Importantly, in the bounded geometry of a cell, broken detailed balance due to the dephosphorylation-phosphorylation cycle implies that a uniform well-mixed state can no longer be a steady state of the system^24^. Instead, all steady states show cytosolic gradients with a density profile that is spatially non-uniform but unpolarised^24^. As the reactive dynamics in the PAR system is bistable, there are two such unpolarised states, one with aPAR and the other with pPAR being the more abundant membrane species. In the zygote, aPARs predominate on the membrane, and we refer to this aPAR-dominant state as the unpolarised state.

To perform a linear stability analysis with respect to this unpolarised state, we use Fourier modes specific for elliptical geometry^23^. These modes are classified as even and odd by their symmetry with respect to reflections through a plane along the long axis, and correspond to patterns aligned along the long and short axes, respectively (Fig. 3*A*). If the real parts of the growth rates *σ* of all modes are negative, small spatial perturbations of the unpolarised state will decay and it will remain stable. In contrast, a positive real part of any growth rate (*σ* > 0) indicates that the unpolarised state is unstable, and initially a pattern will emerge corresponding to the mode with the highest growth rate (Fig. 3*B*). Hence, linear stability analysis can identify the parameter regime where patterns of a certain symmetry (short-vs. long-axis) form spontaneously. On very general grounds ^26, 38^, we expect that bifurcations in mass-conserving reaction-diffusion systems are subcritical and hence these pattern attractors persist over some range outside the linear unstable parameter regime (see also details on FEM simulations in the Method section), where patterns do not form spontaneously but can be triggered by a finite perturbation – such as the fertilisation event.

**Figure 3:**
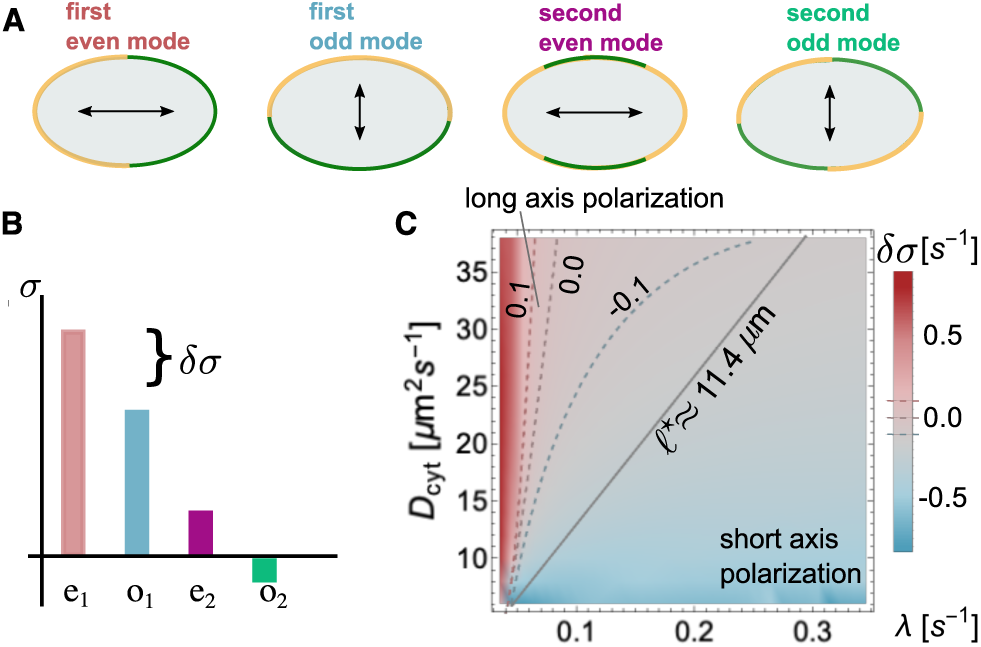
Mode selection and polarity. (*A*) Illustration of the protein distribution on the membrane and the ensuing polarity axis for the lowest-order even and odd modes. (*B*) Illustration of the mode spectrum for these lowest-order modes and the gap *δσ* in the growth rates between the first even and odd modes. (*C*) Relative difference in the growth rates of the first even and odd modes (linear stability analysis in colour code with dashed threshold lines *δσ* = 0*s*^−1^, *δσ* = ±0.1*s*^−1^), *δσ*, as a function of *D*_cyt_ and λ. For small λ and large *D*_cyt_, *δσ* is clearly greater than zero (red, long-axis polarisation), whereas for large λ and small *D*_cyt_, *δσ* lies below zero (blue, short-axis polarisation). These findings are validated using FEM simulations. FEM sweeps in *D*_cyt_ and λ were run until the steady state was reached. These simulations yielded a straight-line interface (black-solid line in (*C*)) in the λ-*D*_cyt_ parameter space which divides long- (above) from short-axis (below) polarisation in steady state. The line corresponds to a constant threshold reactivation length *ℓ*^*^. All other parameters can be found in Table 1.

For a typical cell size and cytosolic diffusion constants in the range of *D*_cyt_ = 5 − 50 *µm*^2^*s*^−1^, linear stability analysis shows that second- and higher-order modes are negligible compared to the first even and odd modes, *σ*_e_ and *σ*_o_. In the parameter regime under consideration, those two growth rates exhibit similar magnitude and at least one of them is positive. To quantify the competition between the first even and odd modes (long-vs. short-axis), we define the relative difference in their growth rates,

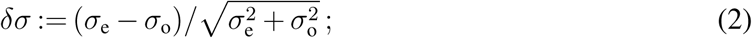

for an illustration see Fig. 3*B*.

### Cytosolic reactivation length is crucial for axis selection

We computed *δσ* as a function of λ and *D*_cyt_. As shown in Fig. 3*C*, the even mode dominates (*δσ* > 0) for large cytosolic diffusion constant and low reactivation rates (favouring long-axis polarisation), otherwise the odd mode dominates. This is consistent with the above heuristic reasoning suggesting that reactivation must be slow or cytosolic diffusion must be fast for the establishment of long-axis polarity. While linear stability analysis can elucidate the selection of the polarisation axis during the onset of pattern formation, it can not predict the final pattern as it neglects nonlinear effects in the diffusion-reaction equation. To determine the final stable polarisation axis we performed finite-element (FEM) simulations; see alslo details on FEM simulations in the Method Section. These simulations show that there is a threshold value for the reactivation length *ℓ*^*^ = 11.4 *µm* above/below which cells stably polarise along the long/short-axis (Fig. 3*C*). We conclude that in a two-dimensional cell geometry the reactivation length *ℓ*, which determines the spatial distribution of active proteins, is the decisive parameter that determines both initial axis selection and its long-term maintenance. How in full three-dimensional cell geometry this effect of the membrane-to-bulk ratio interacts with the role of the interface length will be discussed below.

### Role of phosphorylation rates

Whether there is a spatial separation between aPAR and pPAR domains, is known to depend on the relative magnitude of the phosphorylation rates *k*_Ap_ and *k*_Pa_^9, 16^: an interface between different domains exists and can be maintained only if these antagonistic phosphorylation processes are balanced. To determine the necessary conditions for this balance, we analysed the stability of the unpolarised state using linear stability analysis varying both phosphorylation rates over one order of magnitude. We fixed *D*_cyt_ = 30 *µm*^2^*s*^−1^ and chose two representative reactivation rates, λ = 0.3 *s*^−1^ and λ = 0.05 *s*^−1^, corresponding to reactivation lengths, *ℓ* = 10 *µm* and *ℓ* = 24.5 *µm*, respectively.

Our analysis in elliptical cell geometry shows that spontaneous polarisation starting from the unpolarised state arises only within a limited range of *k*_Pa_*/k*_Ap_ values (cones in Fig. 4), in accordance with previous studies using a one-dimensional model^9, 28^. Strikingly, however, we find that the selection of the polarisation axis does not depend on the mutual antagonism but primarily on the activation-deactivation cycle. The ratio of the phosphorylation rates mainly determines the initial preference for a polarisation axis starting from an unpolarised state (Fig. 4*A* and *B*). Specifically, we find that for λ = 0.3 *s*^−1^, the first even mode grows more slowly than the first odd mode (*δσ* < 0), favouring short-axis polarisation. In contrast, for slower reactivation λ = 0.05 *s*^−1^, the first even mode grows faster than the first odd mode (*δσ* > 0). These respective preferences are most pronounced for large *k*_Pa_*/k*_Ap_. For the mid to low range of *k*_Pa_*/k*_Ap_, one finds *δσ* ≈ 0, i.e. linear stability analysis does not predict a clear preference for either long- or short-axis polarisation. FEM simulations (for details on the FEM simulations see Method Section) show, however, that – irrespective of the ratio *k*_Pa_*/k*_Ap_ – long- and short-axis polarisation in the final steady state is obtained for *ℓ* = 10 *µm* and *ℓ* = 24.5 *µm*, respectively; see Supplementary Movies **M2d 1** – **M2d 3** and Supplementary Tables 2, 3. These simulations confirm that the reactivation length *ℓ* is the deciding factor for axis selection in elliptical geometry.

**Figure 4:**
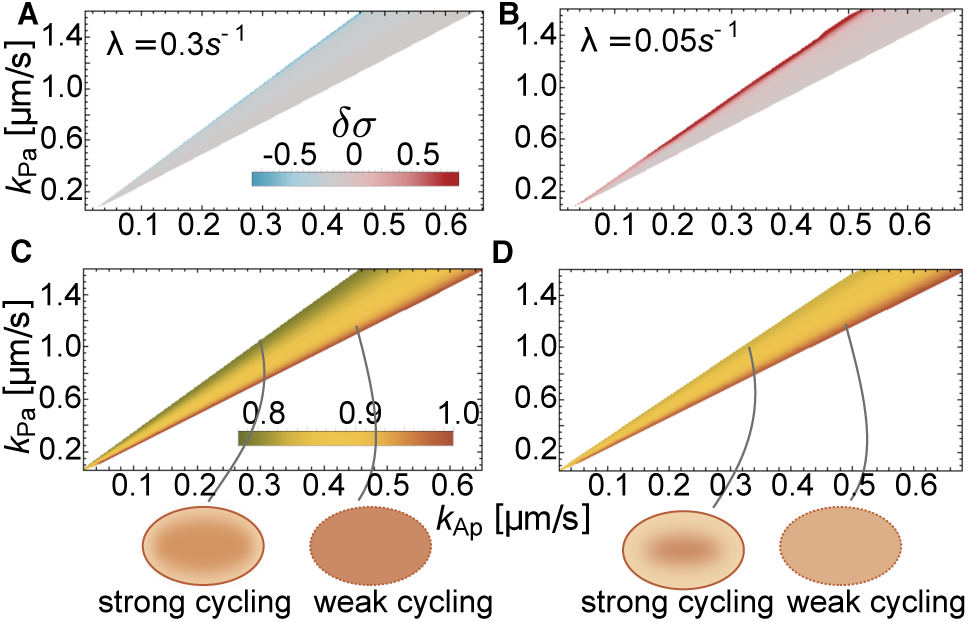
Role of phosphorylation rates in polarisation and axis selection. Linear stability analysis shows that spontaneous polarisation is possible only within a range of ratios of the phosphorylation rates, *k*_Pa_*/k*_Ap_ (cone-shaped regions): The relative difference in the growth rates of even and odd modes (*δσ*) is shown in (*A*) for λ = 0.3 *s*^−1^, and (*B*) for λ = 0.05 *s*^−1^ in colour code (indicated in the graph). Panels (*C*) and (*D*) show the corresponding cytosolic concentration of *A*_1_ in the aPAR dominant unpolarised state (*A*_2_ has a quantitatively similar concentration gradient to *A*_1_ within the cone, not shown), normalised with respect to the maximal concentration of *A*_1_ obtained within the respective cone. Cartoons at the bottom of the figure schematically depict the cytosolic distribution of aPARs throughout the cell.

The FEM simulations further show that outside of the parameter regime of linear instability there exist stable polarised states, showing that the system is excitable, i.e. that patterns can be triggered by a large enough finite perturbation; see Supplementary Notes 1. This parameter regime is actually quite broad (see also Supplementary Fig. 1). As a generic example for an external stimulus, we have investigated how the PAR system reacts to initial concentration gradients on the membrane that were aligned along the final stable polarisation axes. We find that large enough gradients can indeed stimulate the formation of cell polarisation. It would be interesting to specify external cues more in detail experimentally and study how they affect pattern formation. In another work we recently showed that Turing instabilities and excitability (i.e. the ability to establish a pattern by applying a larger perturbation to the stable uniform steady state) are mechanistically linked in mass-conserving systems such as the PAR system ^38^. Hence, even in systems where polarity is established by an external cue, identifying a Turing instability also locates regions where external stimulation leads to stable pattern formation.

The dependence of initial growth rates on the ratio of phosphorylation rates can be attributed to the fact that, in the unpolarised (aPAR-dominant state), the cytosolic concentration of aPARs increases with the rate at which aPARs are phosphorylated by pPARs, i.e. with a reduction in *k*_Pa_*/k*_Ap_ (Fig. 4*C, D*). If a protein species is abundant in the cytosol, recycling of recently detached proteins can be compensated for by a protein of the same type in the cytosolic reservoir attaching to the membrane. Hence, effects due to different membrane-to-bulk ratios in the initial polarisation phase are dominant if the cytosolic pool of proteins undergoing an activation-deactivation cycle is low, explaining why *δσ* depends on geometry for large values of *k*_Pa_*/k*_Ap_ (Fig. 4*C, D*).

### Axis selection depends on relative protein densities

After learning that the abundance of cytosolic proteins determines initial axis selection, we asked how changing the relative total protein densities affects cell polarisation. For all investigations up to this point the average densities were fixed to the order of magnitude determined experimentally by Gross et al.^**?**^ (see Table 1 and see Supplementary Note 2). A linear stability analysis revealed that density variations alter several features: the range of ratios *k*_Pa_*/k*_Ap_ for which an interface between different PAR domains can be stably maintained, and the threshold value of reactivation length *ℓ*^*^ that distinguishes between short- and long-axis polarisation. The effects were most prominent when the ratio of pPAR and aPAR proteins that phosphorylate each other ([*P*]*/*[*A*_2_]), and the ratio of aPAR proteins ([*A*_1_]*/*[*A*_2_]) was varied.

**Table 1:**
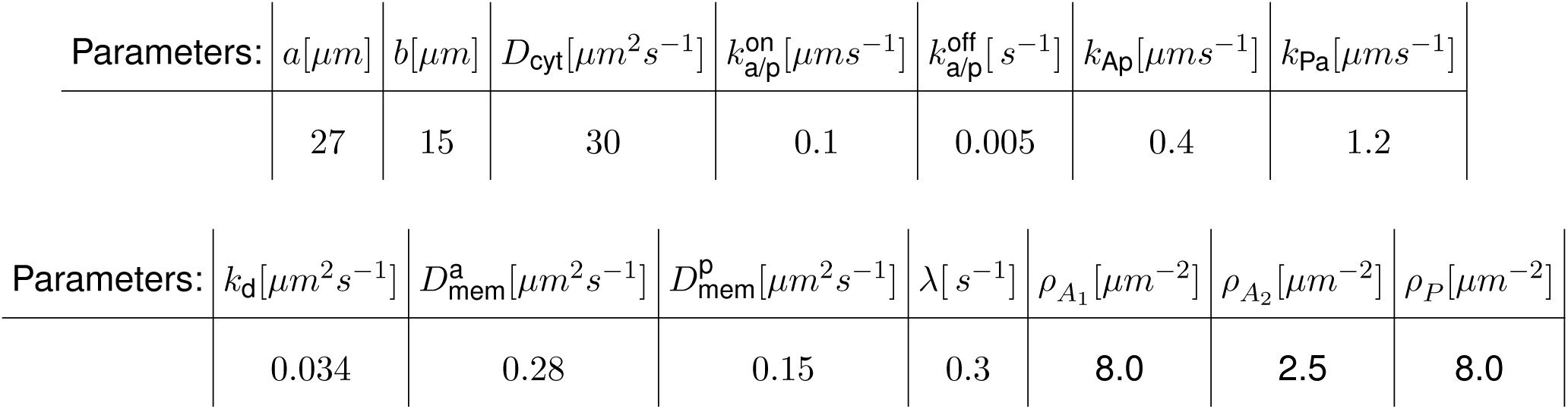
Parameters used to create Fig. 3-5.

As shown in Fig. 5, increasing the ratio of the antagonistic proteins ([*P*]*/*[*A*_2_]) mainly shifts the regime of spontaneous cell polarisation up on the *k*_Pa_*/k*_Ap_ axis. This upward shift is easily explained, as the effective mutual phosphorylation rates are given by *k*_*Ap*_[*P*] and *k*_*Pa*_[*A*_12_], respectively – where [*A*_12_] is mainly limited by the availability of [*A*_2_]. Therefore, when the concentration of pPAR proteins ([*P*]) is increased relative to [*A*_2_], the per capita rate *k*_*Pa*_ has to be increased relative to *k*_*Ap*_ as well, in order to retain the balance between the mutual phosphorylation processes.

**Figure 5:**
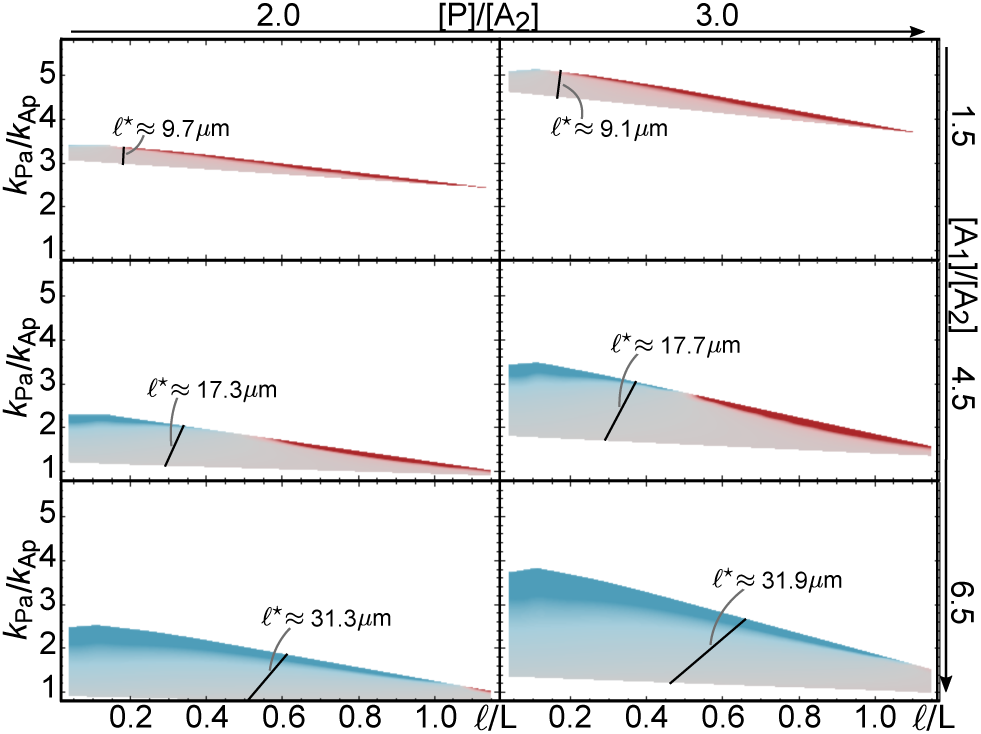
Relative protein numbers determine robustness of cell polarity. Linear stability analysis for a range of density ratios [*P*]*/*[*A*_2_] and [*A*_1_]*/*[*A*_2_]; [*A*_2_] was kept constant. Each graph shows the range of phosphorylation ratios (*k*_Pa_*/k*_Ap_) and relative reactivation lengths (*ℓ/*L) where the base state is linearly unstable, with *δσ* given by the same colour code as in Fig. 4*A*; fixed parameters are *k*_Ap_ = 0.4 *µ*m *s*^−1^ and *D*_cyt_ = 30 *µm*^2^*s*^−1^, and further parameters not varied can be found in Table 1. FEM parameter sweeps of *k*_Pa_ and λ, with fixed parameters *k*_Ap_ = 0.4 *µ*m *s*^−1^ and *D*_cyt_ = 30 *µm*^2^*s*^−1^, for each density set show that the steady state polarisation axis also depends strongly on the ratio [*A*_1_]*/*[*A*_2_]. The steady state switches from short- to long-axis polarisation at the black line in each graph, indicating *ℓ*^*^.

Changing the ratio between the different types of aPAR proteins has two effects. First, spontaneous polarisation is possible for a broader range of *k*_Pa_*/k*_Ap_. Increasing the concentration of the scaffold protein [*A*_1_] relative to [*A*_2_], which phosphorylates pPARs, decreases the lower bound of *k*_Pa_*/k*_Ap_ that allows for polarisation. This is a consequence of the increased reservoir size of *A*_1_ which implies a higher rate of attachment of cytosolic *A*_1_ to the membrane and hence a fast local redimerisation of *A*_2_ (which lacks an inactive phase) right after the detachment of a hetero-dimer *A*_12_. This newly formed hetero-dimer *A*_12_ is then competent to phosphorylate pPARs. Thus it is plausible that even for low *k*_Pa_*/k*_Ap_ one can achieve a balance of mutual antagonism, extending the lower bound of the polarisation regime. Second, changing the ratio [*A*_1_]*/*[*A*_2_] also has a major effect on the threshold value of the reactivation length *ℓ*^*^. We find that *ℓ*^*^ increases with increasing concentration of the scaffold protein [*A*_1_] (Fig. 5). Again, this can be understood as a reservoir effect: globally abundant *A*_1_ promotes immediate re-dimerisation of *A*_2_ with any available *A*_1_. Axis selection is then affected by the polar recycling of *A*_2_.

Taken together, both of these findings emphasise the importance of the activation-deactivation cycle. A cell polarises more robustly when amounts of scaffold proteins are higher. However, at the same time, the cytosolic reactivation length has to increase significantly in order to also robustly maintain long-axis polarisation.

### Role of interface length in three-dimensional cell geometry

With the previous analysis in two-dimensional cell geometry we have built up a basic understanding of the role of the membrane-to-bulk ratio for the selection of the polarisation axis. In a nutshell, we concluded that sufficiently fast diffusion and a sufficiently long inactive phase of the antagonistic proteins ensure that long-axis polarisation is established in a self-organised manner from homogeneous initial membrane concentrations. As the main parameter serving as a proxy for this effect we identified the reactivation length *ℓ*. Is this result directly transferable to a full three-dimensional cell geometry?

Since sensing of the local membrane-to-bulk ratio does not depend significantly on spatial dimension (see also Supplementary Note 4), one would at first sight expect the same conclusions to hold. However, there is a fundamental difference between a three- and a two-dimensional cell geometry. While for an ellipse the interface is always point-like, for a prolate spheroid the interface is longer for short-axis polarisation than for long-axis polarisation; in our case, we have 135 *µm* and 94 *µm*, respectively (Fig. 1 D). This inherent difference between a two- and a three--dimensional cell geometry could significantly affect the protein dynamics on the membrane and in the cytosol. In the absence of an interface the only geometric effect is the membrane-to-bulk ratio. Therefore, as in the two-dimensional case, we expect this ratio to be the main factor that determines the initial formation of the protein domains and the interface between them. However, as soon as an interface has formed, its length is likely to affect the stability of the polarisation axis. The maintenance of the interface between protein domains is presumably energetically costly (protein fluxes sustaining antagonistic reactions, reactivation and rebinding have to be maintained). Therefore, since the interface is longer for short-axis than for long-axis polarisation, it is possible that even an initially favoured alignment of polarisation with the short-axis can become unstable.

To assess the protein dynamics of the system in full cell geometry we performed extensive FEM simulations, restricting ourselves to parameter regimes that we identified as most relevant from the two-dimensional geometry (see Table 4 and compare with Table 1). Starting from a weakly perturbed unpolarised state we observe the following time evolution (Fig. 6 A,B); see the Method section on FEM simulations for 3d system and see our Supplementary Movies 4 and 5 (M3d 1.mp4 and M3d 2.mp4). During an initial time period *T*_initial_ a protein pattern forms that is either aligned along the short or long cell axis or somewhere in between. While long-axis polarisation is stable, any other polarisation is only metastable and after some persistence time *T*_pers_ transitions into stable long-axis polarisation during *T*_trans_; as discussed in Supplementary Note 5 and Supplementary Figures 4-6 there are (unphysiological) cell geometries where short-axis polarisation is stable.

**Figure 6:**
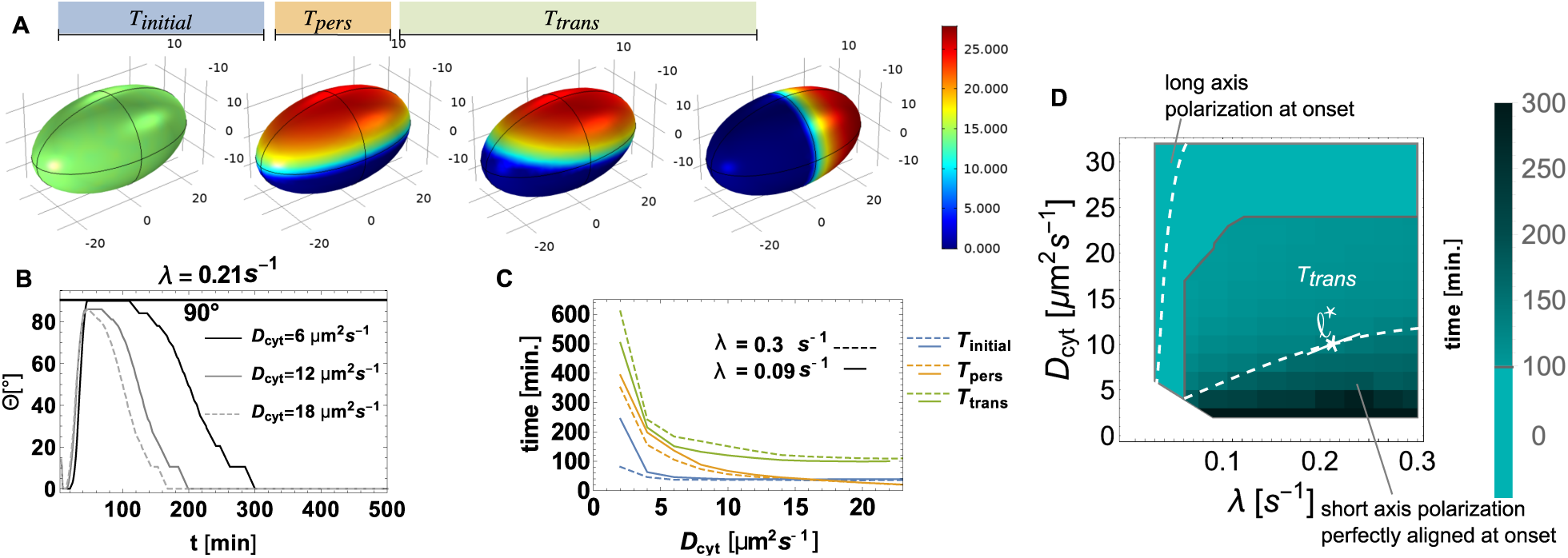
Cell polarisation in three dimensions. (*A*) Image series from FEM (Comsol) simulation for *D*_cyt_ = 6 *µm*^2^*s*^−1^, and reactivation rate λ = 0.21 *s*^−1^. The series illustrates the different times which are further analysed: The time from the initial aPAR-dominated unpolarised state to the initial short-axis polarisation, *T*_initial_; the time duration of persistent short-axis polarisation, *T*_pers_; and the time the pattern takes to turn from short- to long-axis polarisation, *T*_trans_. (*B*) The angle Θ of the concentration maximum of membrane-bound *A*_1_ is plotted against simulation time for different *D*_cyt_ indicated in the graph. (*C*) *T*_initial_, *T*_pers_, *T*_trans_ plotted as a function of *D*_cyt_ for λ = 0.09 *s*^−1^ and λ = 0.3 *s*^−1^. (*D*) The magnitude of the transition time from short- to long-axis polarisation, *T*_trans_, in *D*_cyt_-λ parameter space; a cell was considered to be polarised along the short axis if 90°−10° ≤ Θ ≤ 90°+10°. The monochrome cyan-coloured region above the gray line corresponds to a parameter region where there is no short-axis polarisation, but the polarisation axis is aligned along the diagonal or long axis from the beginning. The dashed lines demarcate parameter regimes where the initial polarisation is aligned perfectly with the short axis (Θ = 90°) or with the long-axis, as indicated in the graph.

We observe that, as for the two-dimensional case, initial long-axis polarisation is favoured for large cytosolic diffusion constants *D*_cyt_ and low reactivation rates λ, while initial short-axis polarisation is favoured for the diametrically opposed case; compare Fig. 6 D with Fig. 3 D. This shows that the local membrane-to-bulk ratio is indeed the main factor that determines initial axis selection. Moreover, the persistence time *T*_pers_ (Fig. 6 C) and the transition time *T*_trans_ (Fig. 6 C,D) both depend strongly on *D*_cyt_ but only weakly on λ. In the regime with a clear preference for short-axis polarisation (below the dashed line in Fig. 6 D), *T*_trans_ becomes as large as several hours; for reference see Fig. 6 D with *ℓ*^*^ ≈ 7*µm*; for further discussion and results on time scales see also Supplementary Note 8 and Supplementary Figure 8.

Finally, we wanted to investigate the main factors that determine the stability of long-versus short-axis polarisation. As the essential novel feature of a three-dimensional cell geometry is the length of the interface between the PAR domains, we speculated that an additional mechanism relevant for axis polarisation is the minimisation of the interface length. To test this hypothesis, we performed FEM simulations in different prolate and oblate geometries; see Supplementary Notes 5 and Supplementary Figures 4 to 6, and Supplementary Movies 6 to 8 (M3d 3 to M3d 5). We find that (for a given set of model parameters) the local diffusive protein fluxes from the cytosol to the membrane at the aPAR-pPAR interface are the same for short- and long-axis polarisation. Hence, the corresponding total fluxes scale with the length of the interface (see also Supplementary Note 6). This suggests that the mechanism responsible for long-axis stability is minimisation of protein fluxes. As a consequence, the transition times *T*_trans_ from short-to long-axis polarisation should also decrease with larger cytosolic protein fluxes as the maintenance of a larger interfaces becomes more costly. Indeed, FEM simulations show that changing the cytosolic diffusion constant leads to an increase in the associated cytosolic fluxes (see Supplementary Note 7 and Supplementary Figs. 7 and 8), and concomitantly to a significant decrease in the transition times *T*_trans_ (Fig. 6 C,D). Taken together, this shows that it is the interplay between membrane-to-bulk ratio and interface length minimisation due to flux (energy) minimisation that drives the selection of the polarisation axis and determines stability and robustness of this selection process.

## Discussion

Here, we have addressed two linked questions concerning cell polarity in *C. elegans*: Under what conditions do cells polarise, and what determines the polarisation axis?

Polarisation in *C. elegans* is controlled by several mechanisms and their interplay: an initial polarisation cue of the centrosome, contraction of the actomyosin network and the PAR reaction-diffusion system which leads to polarisation in a self-organised manner but also interacts with the centrosome as well as with the actomyosin network. Recent research has further revealed some redundant pathways for the reaction-diffusion system depending on other proteins such as CHIN-1, LGL-1 and Cdc42 ^11, 17, 39, 40^. In view of this complexity, it is constructive to disentangle all individual building blocks, mechanical as well as kinetic, and investigate each separately in order to properly identify the underlying mechanisms which (i) leads to polarisation and (ii) aligns it with the long axis. With our work we could now shed light on polarisation and its alignment by the PAR reaction-diffusion system in 2d and in 3d. We expect the insights gained to be essential elements for a future three-dimensional model which combines the reaction-diffusion system with mechanical effects to quantitatively understand pattern formation in the *C. elegans* embryo.

Previous experiments supported by mathematical models in simplified cell geometry have indicated that balance between mutual phosphorylation of aPAR and pPAR proteins is a key mechanism responsible for cell polarisation ^9, 15, 16, 41^. Our theoretical results in realistic cell geometry support this finding. In addition, we have shown that robustness of cell polarity to variations in the phosphorylation rates increases if the scaffold protein PAR-3 is more abundant than PKC-3, which phosphorylates pPARs. Hence, low scaffold abundance is incompatible with robust biological function. This agrees with experimental findings that the scaffold function of PAR-3 is at least partially supported by other proteins (e.g. Cdc-42 ^33^). Our results suggest that it would be worthwhile to experimentally search for other scaffold proteins and test their functional roles in axis selection.

Most importantly, our theoretical analysis in realistic cell geometry reveals that the key processes for axis selection are cytosolic, specifically the cytosolic diffusion and an inactive (phosphorylated) phase of PAR-3 and PAR-2 after detachment from the membrane. The reactivation time (λ^−1^) implies a cytosolic reactivation length 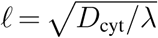 which defines a cytosolic zone of inactive proteins close to the membrane. Proteins with a short reactivation length remain partially caged at the cell poles after membrane detachment, while those with a large reactivation length are uncaged and thereby become uniformly distributed in the cytosol before rebinding. Similarly, proteins lacking a delay, like the PAR-6 PKC-3 complex, are available for rebinding immediately after detachment from the membrane and are thus strongly caged to the cell poles.

Our theoretical analysis in a two-dimensional elliptical geometry shows that only for a sufficiently large cytosolic reactivation length *ℓ* does the long axis become the preferred polarisation axis, at onset as well as for the steady state. For the onset of polarisation, starting from a spatially homogeneous protein distribution, this result is fully transferable to a three-dimensional prolate spheroid. However, in such a realistic cell geometry, the length of the aPAR-pPAR interface also becomes important for the stability of the polarisation axis. Our simulation results suggest an (approximate) extremal principle: The dynamics tries to minimise the interface length such that for physiologically relevant geometries the long axis is always stable. Initial metastable short-axis polarisation is observed if the reactivation length *ℓ* is small (fast reactivation) such that proteins exhibit caging at the polar zones. In that regime, the transition times from short-axis to long-axis polarisation can be of the order of several hours. In contrast, if *ℓ/L* ⪆ 0.3 this time can be as short as 10min. This implies that without guiding cues the reaction-diffusion system requires a sufficiently slow phosphorylation-dephosphorylation cycle and a sufficiently large diffusion constant for fast and robust long-axis polarisation.

Furthermore, how slow reactivation and how fast cytosolic diffusion need to be in order to efficiently and robustly establish and maintain long-axis polarisation depends on the ratio of PAR-3 proteins to the PAR-6 PKC-3 complex: a larger cytosolic pool of PAR-3 attenuates the effect of selecting the interface at midplane and at the same time strengthens the tendency of PKC-3 to put the interface at the poles. Hence we predict that increasing the number of PAR-3 should destabilise long-axis polarisation in favour of short-axis polarisation.

On a broader perspective, these results show that selection of a characteristic wavelength for a pattern and selection of a polarity axis are distinct phenomena and are, in general, mediated by different underlying mechanisms. We expect the following findings to be generic for mass-conserved intracellular protein systems: local membrane-to-bulk ratio and the length of interfaces between different protein domains act as geometric cues for protein pattern formation, and an activation-deactivation as well as cytosolic protein reservoirs alter the sensitivity to cell geometry. Identifying the biochemical steps that are most relevant for axis selection in other intracellular pattern forming systems is an important theme for future research.

## Methods

### Model

First we introduce and discuss the mathematical formulation and analysis of the reaction--diffusion model for PAR protein dynamics. To account for a realistic cell geometry we use, similar as in previous studies of the Min system^23^, a two-dimensional elliptical geometry where the boundary of the ellipse (*∂*Ω) represents the membrane and the interior (Ω) represents the cytosol. Attachment-detachment processes are encoded by nonlinear reactive boundary conditions as introduced in Ref.^23^. Protein interactions are assumed to be bimolecular reactions that follow mass--action law kinetics. In the following a species identifies a mass-conserved protein type, whereas a component indicates the subgroup of proteins in a specific state, such as e.g. ‘phosphorlyated’ (‘inactive’) or ‘membrane bound’.

### Cytosolic dynamics

Proteins in the cytosol are all assumed to diffuse with the same diffusion constant, *D*_cyt_ = 30 *µm*^2^*s*^−1^ (see also Table 1). In addition, we consider dephosphorylation (reactivation) of phosphorylated proteins with an activation (dephosphorylation) rate λ = 0.05/s (see also Table 1). The cytosolic concentration of each protein type *X* is denoted by *c*_*X*_ in its active form and by *c*_*X*_* in its inactive form (if applicable). The dynamics of the bulk components are thus given by the following set of reaction-diffusion equations:

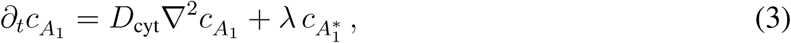

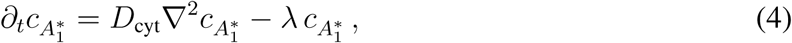

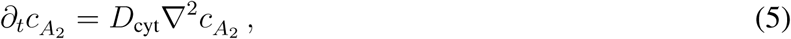

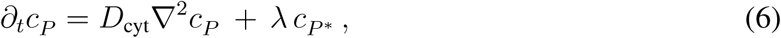

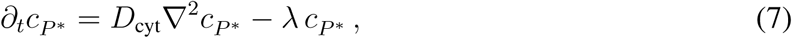

where ∇^2^ is the Laplacian in the two-dimensional bulk.

### Membrane dynamics

On the membrane all species are assumed to diffuse with the respective diffusion constant, 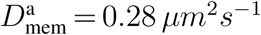 and 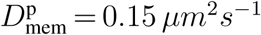 for aPARs and pPARs (see also Table 1). With *m*_*X*_ we denote the membrane-bound concentration of protein *X*. Then, the bimolecular reactions discussed above (see Fig. 1) translate into the following set of reaction-diffusion equations:

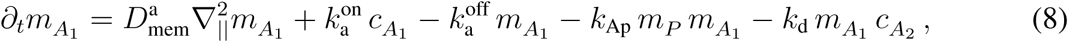

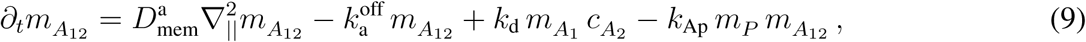

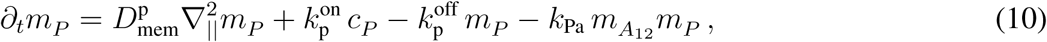

where 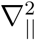 is the Laplacian operator on the boundary *∂*Ω, i.e. on the membrane.

### Reactive boundary conditions

The membrane dynamics and cytosolic dynamics are coupled through reactive boundary conditions. These describe the balance between diffusive fluxes (*D*_cyt_∇_⊥_acting on cytosolic concentration) and attachment and detachment processes between membrane and cytosol:

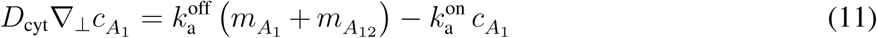

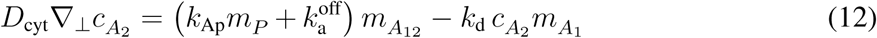

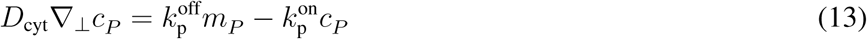

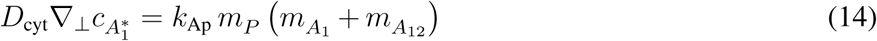

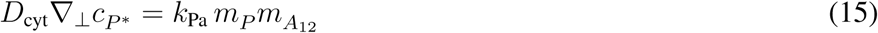

where ∇_⊥_ is the Nabla operator perpendicular to the boundary, such that *D*_cyt_∇_⊥_ is the flux operator between cytosol and membrane.

### Mass conservation

On the time scale of establishment and maintenance of polarisation in *C. elegans*, PAR protein production and degradation are negligible. Hence, the total number *N*_*X*_ of each protein species *X* ∈ {*A*_1_, *A*_2_, *P*} is conserved. It can be obtained by integrating the average densities over the whole space or by integrating the space-dependent cytoplasmic concentrations and membrane concentrations over Ω and *∂*Ω, respectively:

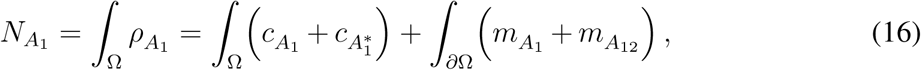

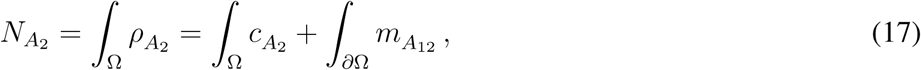

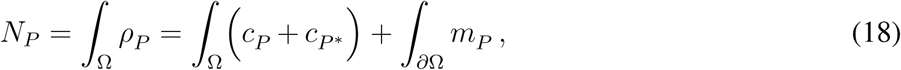

where *∫*_Ω_ and *∫*_*∂*Ω_ denote integrals over the interior and the boundary of the ellipsoid, respectively.

### Linear Stability Analysis

In the following we outline the main steps required to perform a linear stability analysis (LSA) in elliptical geometry, emphasising the major differences relative to the well known stability analysis in planar system geometries with no bulk-boundary coupling (see e.g. a didactic derivation of linear stability analysis written by Cross and Greenside^42^). A detailed derivation of LSA in elliptical geometry can be found in the Supplementary Information of Halatek et al.^23^.

### Reaction-diffusion equations in elliptical geometry

A LSA yields the initial dynamics of a system perturbed from any of its steady states. In the context of pattern formation in reaction-diffusion systems this is typically a uniform steady state. The eigenfunctions of the linearised system (around the steady state) serve as an orthogonal basis in which any perturbation can be expressed. In planar systems these are simply Fourier modes, e.g. ∼ cos(*qx*) with spatial variable *x* and wavenumber *q*, where *q* is chosen such that boundary conditions are satisfied. The LSA then yields the temporal eigenvalues *σ*_*q*_ (growth rates) for each wavenumber that express exponential growth or decay, and possible oscillation (if the imaginary part 𝔍 [*σ*_*q*_] ≠ 0) of the respective eigenfunction exp(*σ*_*q*_*t*) cos(*qx*). Hence, the main objective is (i) to derive the eigenfunctions for the linearised system in the corresponding geometry, and (ii) to calculate the associated growth rates (real parts ℜ[*σ*_*q*_]), where positive growth rates signify formation of patterns with wavelength ∼ 1*/q*. For reaction-diffusion systems with bulk-boundary coupling in elliptical geometry there are three major complications with this approach.

Due to bulk-boundary coupling, we are faced with two separate sets of reaction–diffusion equations. One set is defined in the bulk and accounts for the dynamics in the cytosol. Here reactions are assumed to be linear (first order kinetics) and typically account for nucleotide exchange or (de-)phosphorylation, Eq. (3) – Eq. (7). The second set is defined on the boundary and accounts for the dynamics on the membrane (or cell cortex) including diffusion and reaction, Eq. (8) – Eq. (10).

The first complication arises as follows: Given orthogonal elliptical coordinates

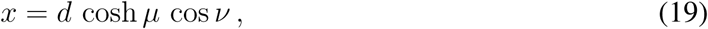

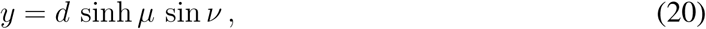

with ‘radial’ variable *µ* > 0, ‘angular’ variable 0 ≤ *ν* < 2*π*, and elliptical eccentricity 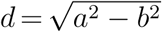 (with long half-axis *a* and short half-axis *b*), the diffusion operator in the bulk *D*_cyt_∇^2^ reads:

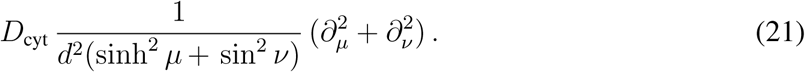

On the boundary the diffusion operator 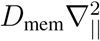 acts along constant *µ* = *µ*_0_ = arctan(*b/a*) and reads:

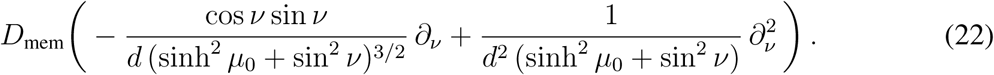

Due to these different diffusion operators the sets of reaction–diffusion equations in the bulk and on the boundary do not share the same set of canonical eigenfunctions (i.e. eigenfunction obtained from separation of variables). To overcome this problem the diffusion on the membrane can be more conveniently expressed in arclength parametrisation *s*(*ν*):

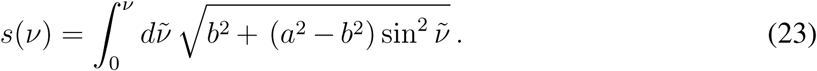

Then, the diffusion operator 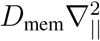 simplifies to 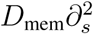, and the eigenfunctions are obtained as

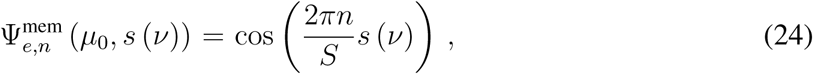

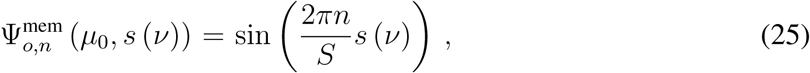

with the circumference of the ellipse *S* = 2*π s*. The goal is then to express these functions in terms of the orthogonal eigenfunctions of the bulk problem — the Mathieu functions, here denoted by Ψ(*ν*) and *R*(*µ*) — which are obtained as solutions of the Mathieu equations:

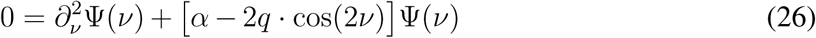

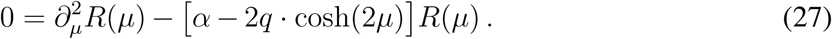

Here *α* is a constant of separation, and

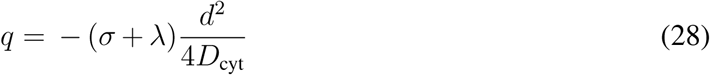

denotes a dimensionless parameter (not to be confused with a wavenumber!). For small *q*, analytical approximations of the Mathieu functions can be obtained ^**?, ?**, 23^ and matched with the eigen-functions 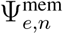 and 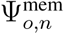 at the boundary *µ* = *µ*_0_.

The second complication is a consequence of the coupling between bulk and boundary processes through the reactive boundary condition, see e.g. the model equations Eq. (11) – Eq. (15). This coupling introduces an explicit dependence of the linearised system on the (derivative of the) radial eigenfunctions *R*(*µ*) (see Ref.^23^), which, in turn, depends on the temporal eigenvalues *σ* in a non-algebraic fashion. Usually, the final step in any LSA is the solution of a characteristic equation 0 = *f* (*σ*), which is typically polynomial in *σ*. Due to the bulk-boundary coupling this is no longer the case (irrespective of the geometry, see e.g. Ref.^26^; the characteristic equation is transcendental and can only be solved numerically for each parameter combination ^23^. Therefore, it is not possible to derive a general stability criterion analogous to that known for planar systems without bulk-boundary coupling ^42^. We further note that the boundary condition introduces a coupling between the angular eigenfunctions Ψ(*ν*), which, however, is small and can be neglected ^23^.

The final complication arrises as consequence of the cytosolic reactivation cycle. This cycle generically precludes the existence of a uniform steady state (including states uniform along the boundary). The origin of this symmetry adaption process has been discussed in Ref.^24^. Following Ref.^23^ we approximate the near-uniform steady state with the eigenfunction that is constant along the boundary, i.e. 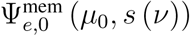. In this case nonlinearities (which are restricted to the boundary) do not induce mode coupling, which would otherwise complicate the LSA.

### Finite Element Simulations (FEM)

Linear stability analysis can only predict the onset of pattern formation. In order to understand the full nonlinear protein dynamics and to determine the steady states corresponding to given parameter sets we further performed finite element (FEM) simulations on a triangular mesh using Comsol Multiphysics 5.1 - 5.4 (updating versions).

### Setup for FEM simulations

As time-dependent solver in Comsol Multiphysics we chose PAR-DISO with a multithreaded nested dissection. The time stepping was performed with a relative tolerance of 10^−6^ between time steps and solved with a multistep method (BDF). In all simulations we used triangular meshing (setting ‘finer’) with additional refinement at the boundary, i.e. along the membrane. As for the linear stability analysis, if not specified otherwise, the parameters for the FEM simulations can be found in Table 1. For the standard parameter sets given in Table 1, we ran the simulation up to 5 · 10^6^*s*. Since the system reached the steady state for most parameter sets at the latest after 5 · 10^5^*s*, we limited simulation times for large parameter sweeps at 10^6^*s*.

### The critical reactivation rate

The 2d FEM sweep of λ versus *D*_cyt_ was initialised with a random initial perturbation of the stationary state with high aPAR concentration on the membrane. The initial perturbation was implemented by drawing a random number rand(*x, y*) from a normal distribution with zero mean and unit variance and multiplying the membrane concentration of aPARs by (1 + 0.01 · rand(*x, y*)) and that of pPARs by (1 − 0.01 · rand(*x, y*)), i.e. we perturbed the initial condition randomly by 1%. The parameter sweep was performed varying λ from 5 · 10^−3^ s^−1^ to 0.3 s^−1^ in steps of 5 · 10^−3^ s^−1^ and varying *D*_cyt_ from 6 *µm*^2^*s*^−1^ to 40 *µm*^2^*s*^−1^ with a uniform spacing of 2 *µm*^2^*s*^−1^.

We further performed two test simulations (sweeping λ and *D*_cyt_) which were initialised with linear gradients. These implementations were intended to uncover dependencies of the final pattern on the initial perturbation. In the first sweep, the gradient was oriented along the long-axis, i.e. the aPAR concentrations were multiplied by (1 + 0.1 · *x/a*) and the pPAR concentrations by (1 − 0.1 · *x/a*). In the second sweep the gradient was oriented along the short-axis, i.e. the aPAR concentrations were multiplied by (1 + 0.1·*y/b*) and the pPAR concentrations by (1−0.1·*y/b*). We found that the steady state polarisation was the same as with small random perturbations. Initial linear gradients with the ‘wrong? alignment only lead to a transient polarisation along the same axis as the initially imposed gradient but then turned to the same polarisation axis as with the random initial perturbation.

Furthermore, we checked the linear stability analysis sweeps on *k*_Ap_ and *k*_Pa_ in Fig. 4 using FEM simulations. The explicit parameter sets *k*_Ap_ and *k*_Pa_ used for probing FEM simulations are shown in Tables 2 and 3. 2d FEM simulations confirm that there λ is the decisive parameter that determines the polarisation axis and not *k*_Ap_ and *k*_Pa_.

**Table 2:** FEM sample sweeps of *k*_*Ap*_, *k*_*Pa*_ with small initial perturbation (1%) for λ = 0.3*s*^−1^.

| $k_{Ap}$ | $k_{Pa}$ | steady state | onset |
| --- | --- | --- | --- |
| 0.44 | 1.68 | no pattern | no pattern |
| 0.46 | 1.62 | no pattern | no pattern |
| 0.48 | 1.56 | no pattern | no pattern |
| 0.5 | 1.5 | short-axis polarisation | short-axis |
| 0.52 | 1.44 | short-axis polarisation | short-axis |
| 0.54 | 1.38 | short-axis polarisation | short-axis |
| 0.56 | 1.32 | short-axis polarisation | long-axis |
| 0.58 | 1.26 | short-axis polarisation | long-axis |
| 0.6 | 1.2 | no pattern | no pattern |

**Table 3:** FEM sample sweeps of *k*_*Ap*_, *k*_*Pa*_ with small initial perturbation (1%) for λ = 0.05*s*^−1^

| $k_{Ap}$ | $k_{Pa}$ | steady state | onset |
| --- | --- | --- | --- |
| 0.44 | 1.68 | no pattern | no pattern |
| 0.46 | 1.62 | no pattern | no pattern |
| 0.48 | 1.56 | no pattern | no pattern |
| 0.5 | 1.5 | long-axis polarisation | long-axis |
| 0.52 | 1.44 | long-axis polarisation | long-axis |
| 0.54 | 1.38 | long-axis polarisation | short-axis |
| 0.56 | 1.32 | long-axis polarisation | short-axis |
| 0.58 | 1.26 | long-axis polarisation | short-axis |
| 0.6 | 1.2 | no pattern | no pattern |

In order to find *ℓ*^*^*/L* in steady state for different combinations of density ratios shown in Fig. 5, we performed FEM sweeps of *k*_Pa_ (for fixed *k*_Ap_ = 0.4 *µms*^−1^) and λ (for fixed *D*_cyt_ = 30 *µm*^2^*s*^−1^) at first in broad steps (the steps for λ were initiated with 5 · 10^−3^ s^−1^ and those for *k*_Pa_ with 0.2 *µms*^−1^). As soon as we identified a regime of parameters for *ℓ*^*^*/L* where long-axis polarisation turned to short-axis polarisation, we used finer steps, with the step size being chosen in accordance with the cone size of each of the *k*_Pa_*/k*_Ap_ versus *ℓ*^*^*/L* cones in Fig. 5.

### FEM simulations for 3d system

In 3d FEM simulations for all sweeps were initiated with an initial *aPAR*-dominant concentration on the membrane and 1% random perturbation thereof. All parameters are shown in Table 4. For the sweep of λ versus *D*_cyt_ resulting in the data discussed in the main text and Fig. 6 the parameter range was set to *D*_cyt_ = 2 − 32 *µm*^2^*s*^−1^ in steps of 2 *µm*^2^*s*^−1^, and reactivation rate λ = 0.03 − 0.3 *s*^−1^ in steps of 0.03 *s*^−1^. The full region of the formation of any pattern can be found by using the feature that the absolute value of membrane gradients is zero for a homogeneous distribution on the membrane and a positive number for inhomogeneous (patterned) protein distributions on the membrane. To distinguish between long and short axis patterns the FEM simulations were analysed by investigating (i) the angle of the concentration maxima on the membrane in ellipsoidal coordinates (which is 90° for perfect short axis polarisation and 0/180° for perfect long axis polarisation) and additionally (ii) the distance between the concentration maximum of *P* and *A*_1_ on the membrane (which is 2·a= 54*µm* for long axis polarisation and 2·b= 30*µm* for short axis polarisation). For a final check, the pattern dynamics was sampled by eye to ensure that these criteria work. In order to numerically investigate the onset of long axis polarisation - which is very sensitive to λ - a finer sweep was additionally performed with *D*_cyt_ = 2 − 32 *µm*^2^*s*^−1^ in steps of 2 *µm*^2^*s*^−1^, and reactivation rate λ = 0.015 − 0.01 *s*^−1^ in steps of *s*^−1^. To find the boundary for a polarity onset with long axis alignment we filtered for a short axis and a diagonal onset.

**Table 4:** Parameters for three-dimensional FEM simulations. For the sweeps shown in Fig. 6 all parameters but λ and *D*_cyt_ were chosen as shown in this Table. λ was varied between 3 · 10^−2^ *s*^−1^ and 0.3 *s*^−1^ with a uniform spacing of 3 · 10^−2^ *s*^−1^. *D*_cyt_ was varied from 2 *µm*^2^*s*^−1^ to 32 *µm*^2^*s*^−1^ with a uniform spacing of 2 *µm*^2^*s*^−1^.

|  |  |  |  |  |  |  |  |
| --- | --- | --- | --- | --- | --- | --- | --- |
| Parameters: | $a[\mu m]$ | $b[\mu m]$ | $D_{\text{cyt}}[\mu m^2 s^{-1}]$ | $k_{\text{a/p}}^{\text{on}}[\mu m s^{-1}]$ | $k_{\text{a/p}}^{\text{off}}[s^{-1}]$ | $k_{\text{Ap}}[\mu m^2 s^{-1}]$ | $k_{\text{Pa}}[\mu m^2 s^{-1}]$ |
|  | 27 | 15 | 30 | 0.1 | 0.005 | 0.4 | 1.2 |
| Parameters: | $k_{\text{d}}[\mu m^3 s^{-1}]$ | $D_{\text{mem}}^{\text{a}}[\mu m^2 s^{-1}]$ | $D_{\text{mem}}^{\text{p}}[\mu m^2 s^{-1}]$ | $\lambda[s^{-1}]$ | $\rho_{A_1}[\mu m^{-3}]$ | $\rho_{A_2}[\mu m^{-3}]$ | $\rho_P[\mu m^{-3}]$ |
|  | 0.034 | 0.28 | 0.15 | 0.3 | 8.0 | 2.5 | 8.0 |

## Data availability

Data supporting the findings of this manuscript are available from the corresponding authors upon reasonable request. A reporting summary for this Article is available as a Supplementary Information file.

## Code availability

Custom written codes used in this study are available from the corresponding author upon reasonable request.

## Acknowledgements

We would like to thank T. Fehm and T. Meinhardt who were involved at early, preliminary stages of this project, and F. Brauns and B. Osberg for critical reading of the manuscript and for providing valuable feedback. E.F. acknowledges support from the Deutsche Forschungsgemeinschaft (DFG) via project B02 within the Collaborative Research Center Deutsche Forschungsgemeinschaft (DFG, German Research Foundation) - Project-ID 201269156 - SFB 1032. R.G. is supported by a DFG fellowship through the Graduate School ‘Quantitative Bioscience Munich’ (QBM).

## Author contributions

R.G., J.H., L.W., and E.F. designed research, performed research, and wrote the paper.

---

**Fig. 3**: For the sweep using linear stability analysis in Fig. 3 **C** all parameters but λ and *D*_cyt_ were chosen as shown in this Table. λ was varied between 5 · 10^−3^ *s*^−1^ and 0.35 *s*^−1^ with a uniform spacing of 5 · 10^−3^ *s*^−1^. *D*_cyt_ was varied from 6 *µm*^2^*s*^−1^ to 38 *µm*^2^*s*^−1^ with a uniform spacing of 2 *µm*^2^*s*^−1^. **Fig. 4**: For the linear stability analysis sweep in Fig. 4 **A**,**B** all parameters but λ and *k*_Ap_ and *k*_Pa_ were chosen as above. *k*_Ap_ was varied between 0.02*µms*^−1^ and 0.8*µms*^−1^ and *k*_Pa_ was varied between 0.06*µms*^−1^ and 1.6*µms*^−1^; for both parameters values were uniformly spaced with distance 0.02*µms*^−1^. **Fig. 5**: For the linear stability analysis sweeps in Fig. 5 all parameters but the densities *ρA*_1_, *ρA*_2_ and *ρP*, λ and *k*_Ap_ were set as shown above. For all triples of densities *ρA*_2_ = [*µm*^−2^] while *ρA*_1_ and *ρP* were varied accordingly. The simultaneous sweep of *ℓ* and *k*_Pa_*/k*_Ap_ was obtained by varying λ and *k*_Pa_ for fixed *D*_cyt_ = 30 *µm*^2^*s*^−1^ and *k*_Ap_ = 0.4*µms*^−1^.The values of *ℓ* were uniformly spaced from 2*µm* to 62*µm* with distance 2*µm*. The ratio *k*_Pa_*/k*_Ap_ was varied from 0.7 to 8.0 with uniform steps of 0.05.

## References

1. Campanale, J. P., Sun, T. Y. & Montell, D. J. Development and dynamics of cell polarity at a glance. J Cell Sci 130, 1201–1207 (2017).

2. Chiou, J.-g., Balasubramanian, M. K. & Lew, D. J. Cell polarity in yeast. Annu Rev Cell Dev Biol 15, 365–391 (2017).

3. Roignot, J., Peng, X. & Mostov, K. Polarity in mammalian epithelial morphogenesis. Cold Spring Harb Protoc 33 (2013).

4. Goehring, N. W. PAR polarity: from complexity to design principles. Exp Cell Res 328, 258–266 (2014).

5. Goldstein, B. & Macara, I. G. The PAR proteins: fundamental players in animal cell polarization. Dev Cell 13, 609–622 (2007).

6. Gönczy, P. Asymmetric cell division and axis formation in the embryo. WormBook 1–20 (2005).

7. Cuenca, A. A., Schetter, A., Aceto, D., Kemphues, K. & Seydoux, G. Polarization of the C. elegans zygote proceeds via distinct establishment and maintenance phases. Development 130, 1255–1265 (2003).

8. Munro, E., Nance, J. & Priess, J. R. Cortical flows powered by asymmetrical contraction transport PAR proteins to establish and maintain anterior-posterior polarity in the early C. elegans embryo. Dev Cell 7, 413–424 (2004).

9. Goehring, N. W. et al. Polarization of PAR proteins by advective triggering of a pattern-forming system. Science 334, 1137–1141 (2011).

10. Motegi, F. et al. Microtubules induce self-organization of polarized PAR domains in Caenorhabditis elegans zygotes. Nat Cell Biol 13, 1361–7 (2011).

11. Motegi, F. & Seydoux, G. The PAR network: redundancy and robustness in a symmetry-breaking system. Philos Trans R Soc Lond B Biol Sci 368, 20130010 (2013).

12. Hill, D. P. & Strome, S. An analysis of the role of microfilaments in the establishment and maintenance of asymmetry in Caenorhabditis elegans zygotes. Dev Biol 125, 75–84 (1988).

13. Cowan, C. R. & Cowan, A. A. Centrosomes direct cell polarity independently of microtubule assembly in C. elegans embryos. Nature 431, 92–96 (2004).

14. Petrásek, Z. et al. Characterization of protein dynamics in asymmetric cell division by scanning fluorescence correlation spectroscopy. Biophys J 95, 5476–5486 (2008).

15. Dawes, A. T. & Munro, E. M. PAR-3 oligomerization may provide an actin-independent mechanism to maintain distinct par protein domains in the early Caenorhabditis elegans embryo. Biophys J 101, 1412–1422 (2011).

16. Goehring, N. W., Hoege, C., Grill, S. W. & Hyman, A. PAR proteins diffuse freely across the anterior-posterior boundary in polarized C. elegans embryos. J Cell Biol 193, 583–594 (2011).

17. Schonegg, S. & Hyman, A. A. Cdc-42 and rho-1 coordinate acto-myosin contractility and par protein localization during polarity establishment in c. elegans embryos. Development 133, 3507–3516 (2006).

18. Schonegg, S., Constantinescu, A. T., Hoege, C. & Hyman, A. A. The Rho GTPase-activating proteins RGA-3 and RGA-4 are required to set the initial size of PAR domains in Caenorhabditis elegans one-cell embryos. Proc Natl Acad Sci USA 104, 14976–14981 (2007).

19. Schenk, C., Bringmann, H., Hyman, A. A. & Cowan, C. R. Cortical domain correction repositions the polarity boundary to match the cytokinesis furrow in C. elegans embryos. Development 137, 1743–1753 (2010).

20. Hoege, C. & Hyman, A. A. Principles of PAR polarity in Caenorhabditis elegans embryos. Nat Rev Mol Cell Biol 14, 315–322 (2013).

21. Hao, Y., Boyd, L. & Seydoux, G. Stabilization of cell polarity by the c. elegans ring protein par-2. Dev Cell 10, 199–208 (2006).

22. Zonies, S., Motegi, F., Hao, Y. & Seydoux, G. Symmetry breaking and polarization of the C. elegans zygote by the polarity protein PAR-2. Development 137, 1669–1677 (2010).

23. Halatek, J. & Frey, E. Highly canalized MinD transfer and MinE sequestration explain the origin of robust MinCDE-protein dynamics. Cell Rep 1, 741–752 (2012).

24. Thalmeier, D., Halatek, J. & Frey, E. Geometry-induced protein pattern formation. Proc Natl Acad Sci USA 113, 548–553 (2016).

25. Wu, F., Halatek, J., Reiter, M., Kingma, E. & Frey, E. Multistability and dynamic transitions of intracellular Min protein patterns. Mol Syst Biol 12, 873 (2016).

26. Halatek, J. & Frey, E. Rethinking pattern formation in reaction-diffusion systems. Nat Phys 14, 507–514 (2018).

27. Dawes, A. T. & Iron, D. Cortical geometry may influence placement of interface between Par protein domains in early Caenorhabditis elegans embryos. J Theor Biol 333, 27–37 (2013).

28. Trong, P. K., Nicola, E. M., Goehring, N. W., Kumar, K. V. & Grill, S. W. Parameter-space topology of models for cell polarity. New J Phys 16, 65009 (2014).

29. Tabuse, Y. et al. Atypical protein kinase C cooperates with PAR-3 to establish embryonic polarity in Caenorhabditis elegans. Development 125, 3607–3614 (1998).

30. Etemad-Moghadam, B., Guo, S. & Kemphues, K. Asymmetrically distributed PAR-3 protein contributes to cell polarity and spindle alignment in early C. elegans embryos. Cell 83, 743– 752 (1995).

31. Watts, J. L. et al. par-6, a gene involved in the establishment of asymmetry in early C. elegans embryos, mediates the asymmetric localization of PAR-3. Development 122, 3133–3140 (1996).

32. Hung, T.-J. & Kemphues, K. J. PAR-6 is a conserved PDZ domain-containing protein that colocalizes with PAR-3 in Caenorhabditis elegans embryos. Development 126, 127–135 (1999).

33. Rodriguez, J. et al. aPKC cycles between functionally distinct PAR protein assemblies to drive cell polarity. Dev Cell 42, 400–415.e9 (2017).

34. Li, J. et al. Binding to PKC-3, but not to PAR-3 or to a conventional PDZ domain ligand, is required for PAR-6 function in C. elegans. Dev Biol 340, 88–98 (2010).

35. Li, B., Kim, H., Beers, M. & Kemphues, K. Different domains of C. elegans PAR-3 are required at different times in development. Dev Biol 344, 745–757 (2010).

36. Boyd, L., Guo, S., Levitan, D., Stinchcomb, D. T. & Kemphues, K. J. PAR-2 is asymmetrically distributed and promotes association of P granules and PAR-1 with the cortex in C. elegans embryos. Development 122, 3075–3084 (1996).

37. Driever, W. & Nüsslein-Volhard, C. A gradient of bicoid protein in Drosophila embryos. Cell 54, 83–93 (1988).

38. Brauns, F., Halatek, J. & Frey, E. Phase-space geometry of reaction–diffusion dynamics. Preprint at 1812.08684 (2018).

39. Sailer, A., Anneken, A., Li, Y., Lee, S. & Munro, E. Dynamic Opposition of Clustered Proteins Stabilizes Cortical Polarity in the C.elegans Zygote. Dev Cell 35, 131–142 (2015).

40. Beatty, A., Morton, D. G. & Kemphues, K. PAR-2, LGL-1 and the CDC-42 GAP CHIN-1 act in distinct pathways to maintain polarity in the C. elegans embryo. Development 140, 2005–2014 (2013).

41. Tostevin, F. & Howard, M. Modeling the establishment of PAR protein polarity in the one-cell C. elegans embryo. Biophys J 95, 4512–4522 (2008).

42. Cross, M. & Greenside, H. Pattern Formation and Dynamics in Nonequilibrium Systems (Cambridge University Press, 2009).

