## Supplementary Document for "Geometric cues stabilise long-axis polarisation of PAR protein patterns in *C. elegans*"

Raphaela Geßele, Jacob Halatek, Laeschkir Würthner & Erwin Frey*
Arnold Sommerfeld Center for Theoretical Physics
and Center for NanoScience, Department of Physics,
Ludwig-Maximilians-Universität München,
Theresienstraße 37, 80333 München, Germany.

### SUPPLEMENTARY NOTES

Following the structure of the main text, Supplementary Note 1 discusses further aspects of cell polarisation in two-dimensional elliptical geometry using finite element simulations: excitable region in parameter space, and time evolution of the polarisation axis. Supplementary Note 2 summarises experimental information on protein numbers. In Supplementary Note 3 we show why it is not sufficient to use a planar geometry in order to learn about the selection of the polarisation axis. The membrane-to-bulk ratio in two-dimensional elliptical and three-dimensional ellipsoidal geometry is summarised in Supplementary Note 4. In order to challenge the hypothesis of interface minimisation, in Supplementary Note 5 the results on axis selection in oblate and prolate geometries are discussed. To understand the relative role of the activation-deactivation cycle and interface minimisation an extensive set of finite element simulations was performed and the results are discussed. Supplementary Note 6 and 7 show that interface minimisation arises from flux minimisation. Finally, in Supplementary Note 8 patterning time scales are provided and discussed.

#### Supplementary Note 1

**Stimulus-induced polarisation and transient polarisation alignment.** In the wild type *C. elegans* embryo polarisation is established by an interplay between mechanical cues (forces of the centrosome after male sperm entry and actomyosin contraction towards the anterior) and the PAR reaction diffusion system. In the main text we focused on spontaneous pattern formation facilitated by a Turing instability. Here, we investigate whether the Turing instability is subcritical, i.e. whether patterns can be induced (stimulated) by large perturbations outside the Turing unstable region, such as the fertilization event. To this end, we performed FEM simulations that were initiated with linear concentration gradients along the membrane as initial conditions. The gradient was chosen to favor selection of a pattern aligned with the same polarisation axis as predicted by linear stability analysis.

Specifically, for $\lambda = 1s^{-1}$ (fast reactivation) shown in Supplementary Figure 1(**A**) top row, the gradient was chosen along the short axis, i.e. the aPAR concentrations were multiplied by $(1 + y/b)$ and the pPAR concentrations by $(1 - y/b)$, where $2b$ is the length of the short axis. For $\lambda = 0.05s^{-1}$ (slow reactivation) shown in Supplementary Figure 1**(B)** top row, the gradient was aligned along the long axis, i.e. the aPAR concentrations was multiplied with $(1 + x/a)$ and the pPAR concentrations with $(1 - x/a)$, where $2a$ is the length of the long axis. Indeed, in both cases we found a large parameter domain outside the regime of spontaneous polarisation where pattern formation can be triggered by finite perturbations. The specific sets $(k_{\mathrm{Ap}}, k_{\mathrm{Pa}})$ for which the system was tested for stimulus-induced pattern formation are provided in the Tables 1, 2 and 3. In Supplementary Figure 1 we extrapolated from this data to find an outer cone of the excitable region (dashed lines in the $k_{\mathrm{Ap}}$-$k_{\mathrm{Pa}}$ diagrams).

Furthermore, in the Turing unstable regime we tested alignment of polarisation when the initial condition in the FEM simulation was chosen to select for the pattern orthogonal to the polarisation axis predicted by linear stability analysis. For all sets of parameters

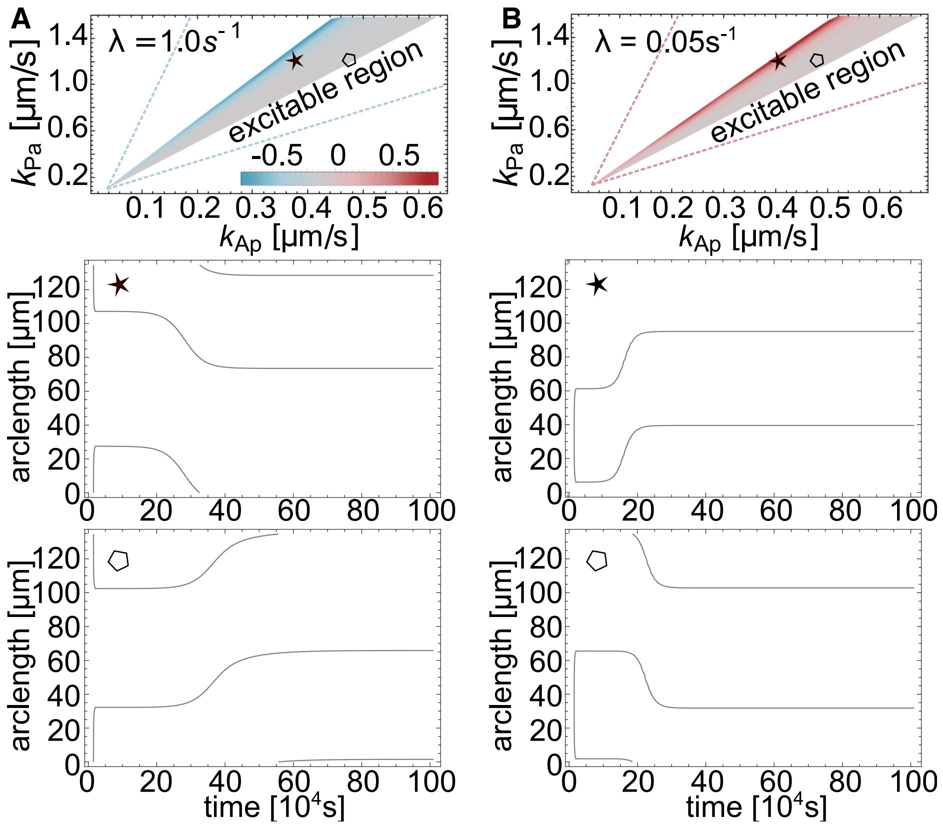


Supplementary Figure 1. Testing the steady state polarity axis with initial gradients. (**A,B**) top row: Investigation of excitable region. The cones show the results of a linear stability analysis in the 2d ellipse as a function of $k_{Ap}$ and $k_{Pa}$ with the color code indicating the normalized difference of the first even and odd growth rate, $\delta\sigma$ (same color code as in Fig. 4 of the main text, i.e. even mode grows faster: red, odd mode grows faster: blue, $\delta\sigma \approx 0s^{-1}$: gray). These Turing-unstable regions are flanked by parameter regimes (bounded by dashed lines), where patterns can only be induced by a large enough stimulus acting on the uniform state; we call this the excitable region. Parameters as in Table 1 of the main text. (**A,B**) Middle and bottom row: Investigation of polarisation re-alignment. The black lines indicate the interface position between aPAR and pPAR domain. Shown are sample interface trajectories from FEM simulations for parameters at the upper (star) and lower (hexagon) bound of the Turing-unstable regime. In contrast to the top row of the Figure, here the FEM simulations were initialised with gradients aligned perpendicularly to the predicted pattern orientation (gradients as above, but orthogonal to predicted polarisation alignment, for mathematical definition see text). We find that the initial polarisation axis is aligned with the initial gradient while the final pattern is dictated by the reactivation cycle. $k_{Ap}$ and $k_{Pa}$ do not impact this qualitatively but only the transition time from one to the other polarisation axis.

$(k_{\mathrm{Ap}}, k_{\mathrm{Pa}})$ which we tested we found that the final steady state was the same as the one predicted by linear stability analysis (at the upper bound of $(k_{\mathrm{Pa}}, k_{\mathrm{Ap}})$ where $\delta\sigma$ is decisively above or below zero). However, a transiently lasting polarisation along the axis of the initial gradient was observed (see Supplementary Figure 1 middle and bottom row). In detail, for fast $\lambda = s^{-1}$ the initial gradient is aligned with the long axis, i.e. the original aPAR concentrations were multiplied with $(1+x/a)$ and the pPAR concentrations with $(1-x/a)$. Polarisation establishes along the long axis first (for both pairs of $(k_{Ap}, k_{Pa})$ in Supplementary Figure 1, "star" and "hexagon"), and then transitions to align with the short axis where it then finds its steady state. This turning of the polarisation axis starts later for lower $k_{Pa}/k_{Ap}$ ratios (compare Supplementary Figure 1, bottom row). For slow $\lambda = 0.05s^{-1}$ we find just the opposite behaviour: Initial short axis polarisation establishes aligned with the gradient but then turns towards steady state long axis polarisation. The time of turning again depends on the ratio $k_{Pa}/k_{Ap}$.

### Supplementary Note 2

**Total and relative protein numbers.** For the PAR protein system in *C. elegans*, many parameters have been measured including relative and total protein numbers, binding and unbinding rates, and diffusion constants of proteins on the membrane [1–4]. However, measurements of the PAR protein density were reported with a relatively large uncertainty; according to the Supplementary Material in Ref. [1] with a relative error larger than 20%. Most recent experiments report total PAR protein densities between 2 and 6 proteins per $\mu m^3$ if all proteins were evenly distributed in the cytosol (depending on the specific PAR protein) [4]. We used the corresponding order of magnitude of total protein numbers (see Table 1 of the main text) for our studies and further investigated relative abundances of proteins (see the relative density variations $[P]/[A_2]$ and $[A_1]/[A_2]$ discussed in Section "Robustness of polarisation as well as axis selection depend on the relative protein densities") and Fig. 5 in the main text).

### Supplementary Note 3

**Planar geometry: the characteristic lengthscale does not select the axis.** Is it possible to simplify the geometry of a cell in order to answer the question of axis selection for cell polarisation? A heuristic argument in favor of a positive answer would be: Let's simplify to a planar geometry as illustrated in Fig 2 A, and perform a linear stability analysis. This will yield a fastest growing mode at some characteristic wavelength. Intuitively, one may now expect that in elliptical geometry those axis is selected which length fits this characteristic wavelength best. Is this intuition correct?

To answer this question, we investigated the PAR model in planar geometry and compared it with the results that we obtained in elliptical and ellipsoidal geometry (main text). The linear stability analysis was performed in a rectangular two-dimensional geometry $(x, z)$ with variable width and fixed height $h$ that matches the short half-axis $b$ of the ellipsoidal cell; see Supplementary Figure 2 A. The membrane is at the bottom, $z = 0$, where we assume reactive boundary conditions. For symmetry reasons we assume no-flux boundary conditions at $z = b$. The details of the linear stability analysis can be found in Ref. [5]. The

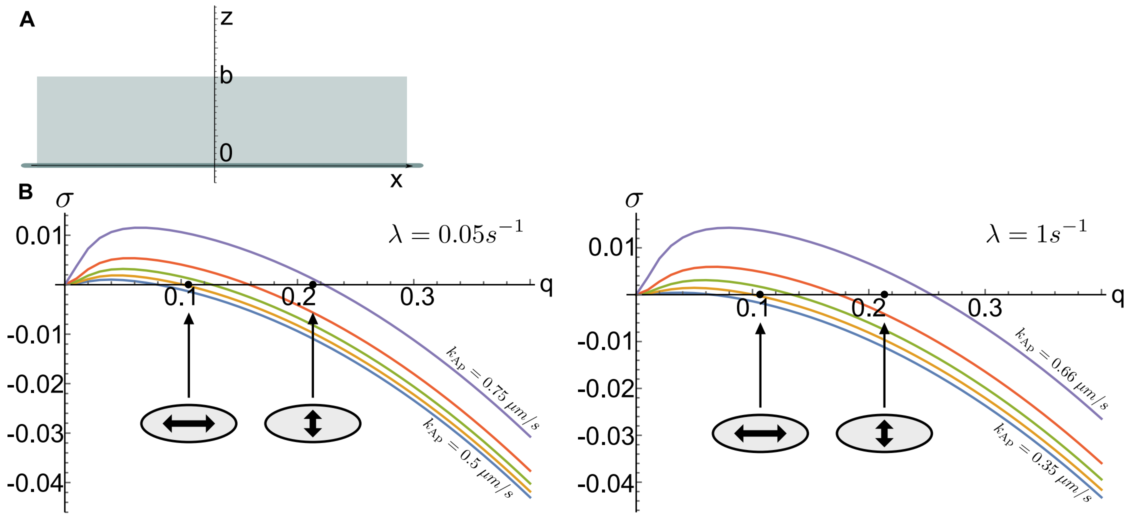


Supplementary Figure 2. Linear stability analysis in planar geometry. (**A**) Illustration of a planar geometry with membrane at the bottom, $z = 0$, and cytosol of height $h = b$. (**B**) Dispersion relations in rectangular geometry for $\lambda = 0.05\,\mathrm{s}^{-1}$ (left, long-axis selection in the ellipse) and $\lambda = 1\,\mathrm{s}^{-1}$ (right, short-axis selection in the ellipse), showing that the fastest growing mode depends sensitively on $k_{\mathrm{Ap}}$. The filled black circles highlight the length scales corresponding to long axis polarity $q = \pi/(2a)$ and short axis polarity $q = \pi/(2b)$. Naively, the stability analysis in rectangular geometry suggests that modes with large length scale (long axis polarity) are always preferred, contradicting the correct results from the simulations and linear stability analysis in elliptical geometry.

numerical values of all parameters are unchanged (i.e. as in Table 1 in the main text), except the attachment rates $k^{\mathrm{on}}_{\mathrm{a/p}}$ which we rescaled to $0.3 \mu m s^{-1}$] to recover the lateral (Turing) instability of the unpolarised aPAR state. A parameter sweep of the phosphorylation rate constant $k_{\mathrm{Ap}}$ for $\lambda = 0.05\,\mathrm{s}^{-1}$ (long axis selection in the ellipse) and $\lambda = 1\,\mathrm{s}^{-1}$ (short axis selection in the ellipse) shows that the band of unstable modes and the *fastest growing mode* (the mode which determines the *characteristic length scale* at onset) sensitively depend on $k_{\mathrm{Ap}}$ but not on $\lambda$. Furthermore, we find that the *fastest growing mode* corresponds to a *characteristic length scale* which is always longer than the short axis of the cell, $2b$, and can be tuned to fit the long axis, $2a$ (see also marks in the dispersion relation in Supplementary Figure 2 B). Following the heuristic argument one would conclude that the long axis is chosen for polarisation because it fits better into the cell. However, our results in the main text demonstrate that axis selection in cellular geometry is determined by cytosolic parameters such as $\lambda$ and $D_{\mathrm{cyt}}$, but effects by $k_{\mathrm{Ap}}$ are negligible. Hence, we conclude that the *characteristic length scale* determined by linear stability in planar geometry does neither inform about axis selection in elliptical nor ellipsoidal geometry.

We find that pattern alignment is a process which strongly depends on the distribution of binding active proteins in the cytosol. For our model pattern alignment is not dictated by the wavelength of a pattern but rather by the reactivation length $\ell^*$ and the topology of the domain interfaces.

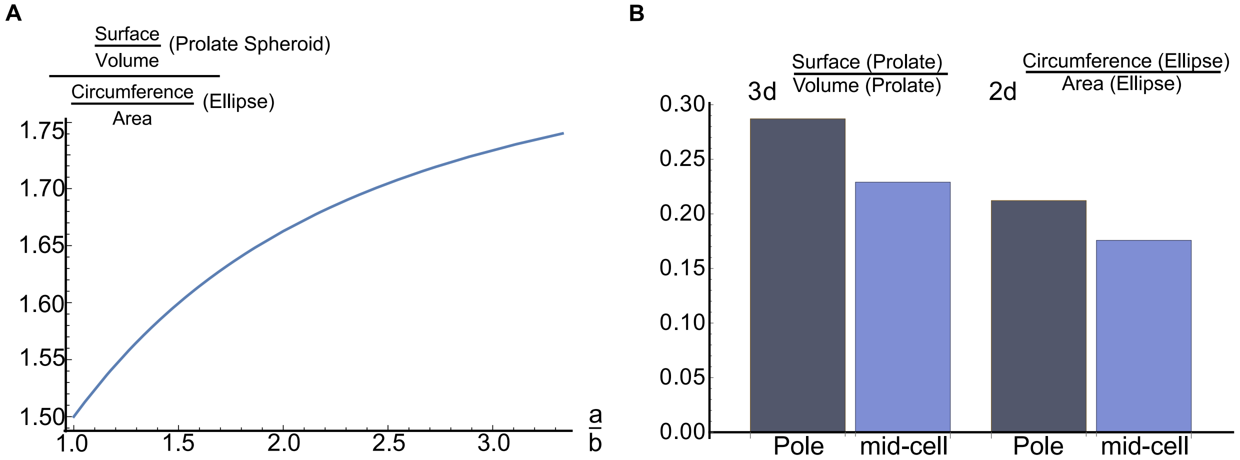


Supplementary Figure 3. Membrane-to-bulk ratio for a two-dimensional (2d) ellipse and a three--dimensional (3d) prolate spheroid. (**A**) The *overall* membrane-to-bulk ratio (integrated over the whole cell boundary) of a prolate spheroid is compared to that of an ellipse with the same minor and major axes as a function of the aspect ratio $a/b$. (**B**) The *local* membrane-to-bulk ratio (as defined in the main text of the supplement) of a prolate spheroid and an ellipse, both at the cell poles and at midcell. The membrane-to-bulk ratio was calculated for some sample diffusion length $\ell_D = 7.5\mu m$.

### Supplementary Note 4

**Membrane-to-bulk ratio for ellipses and prolate spheroids.** In the main text we showed that the membrane-to-bulk ratio is a key factor for axis selection, especially during the initial phase of pattern formation. How does this ratio depend on the dimensionality of the system? Figure 3**A** compares the *overall* membrane-to-bulk ratio — the ratio of area/circumference of the membrane to volume/area of the cytosol ('bulk') — for a two-dimensional ellipse and a three-dimensional prolate spheroid (ellipsoid). One observes that this ratio is in general larger for a prolate spheroid, and the surplus is increasing with the aspect ratio $a/b$.

The *local* membrane-to-bulk ratio varies qualitatively in a similar fashion for the two- and three-dimensional case: it is maximal at the poles and decreases monotonously towards midcell where it reaches its minimum. In order to see this quantitatively we have calculated the membrane-to-bulk ratio for a sample diffusion length ($\ell_D = 7.5\,\mu m$) at the poles and at midcell for an ellipse and a prolate spheroid; see Supplementary Figure 3**B**. To determine the membrane-to-bulk ratio at the poles, we defined a sphere with the sample diffusion length $\ell_D = 7.5\,\mu m$ as radius and center at the cell pole. Then we calculated the membrane region of the ellipse (2d) or ellipsoid (3d) which lies within this sphere. This gives the membrane part of the membrane-to-bulk ratio. The bulk part was calculated by the intersecting region of the ellipse (2d) or ellipsoid (3d) with the sphere. Similarly, we defined the membrane to bulk ratio at midcell with the help of a sphere with radius $\ell_D$ and center at the midcell membrane. We find that quantitatively, the change in the local membrane-to-bulk ratio from midcell to pole is more pronounced in a three-dimensional prolate spheroid than in a two-dimensional ellipse.

### Supplementary Note 5

**The role of interface length for the selection of the polarity axis.** We argued in the main text, that axis selection during cell polarisation is determined by an interplay between two effects: The higher membrane-to-bulk ratio at the cell poles favors short-axis selection for small enough reactivation lengths $\ell$. Otherwise, long-axis polarisation is favored. This is confirmed by our studies for two-dimensional ellipses; see also Method Section *The critical reactivation rate to switch steady state polarity*. On the other hand, we have argued in the main text that there is a tendency of the dynamics to minimize the length of the interface between aPAR and pPAR domains, which would always favor long-axis polarisation.

In this section we give a detailed account of FEM simulations Comsol Multiphysics 5.4 for various three-dimensional ellipsoidal geometries including both prolate and oblate spheroids; see Tables 4, 5. The goal is to clarify the relative role of the membrane-to-bulk ratio and the interface length in the axis selection process.

*a.* ***Perimeter ratio for long- and short-axis polarisation in prolate and oblate spheroid geometries.*** The interface length for short-axis polarisation, $L_{\text{short}}$, and long-axis polarisation, $L_{\text{long}}$ are different for an ellipsoidal geometry. There is a difference between prolate and oblate geometries insofar as the ratio of the interface length for long- and short-axis polarisation, $L_{\text{long}}/L_{\text{short}}$ (short: perimeter ratio), differs; for an illustration see Supplementary Figure 4.

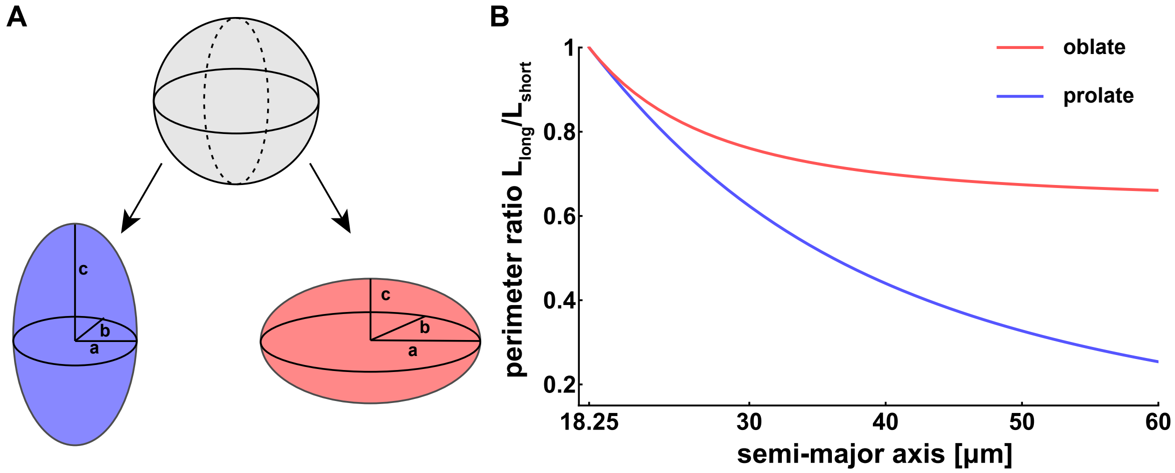


Supplementary Figure 4. Perimeter ratio for an oblate and a prolate spheroid. **(A)** Smooth deformation of a sphere (top) to a prolate spheroid (bottom, left) or to an oblate spheroid (bottom, right) with the same volume as the sphere. Note that for a prolate $a = b < c$ while for an oblate $a = b > c$. **(B)** The perimeter ratio, $L_{\text{long}}/L_{\text{short}}$, as a function of the length of the semi-major axis for an oblate (red curve) and a prolate spheroid (blue curve) with the same volume as a sphere of radius $R = 18.25\,\mu m$

We compare the perimeter ratio $L_{\text{long}}/L_{\text{short}}$ for three-dimensional ellipsoids of the same volume. As our reference system we us a prolate spheroid with axes $15\,\mu m - 15\,\mu m - 27\,\mu m$, i.e. $a = b = 15\mu m$ (semi-minor axis) and $c = 27\,\mu m$ (semi-major axis). This is the same geometry that has been used to generate the results shown in the main text. The volume of an ellipsoid is $V_{\text{ellipsoid}} = \frac{4\pi}{3}a^2c$, corresponding to a sphere of same volume with radius $R = (a^2c)^{1/3} = 18.25\,\mu m$. Note that in contrast to a prolate spheroid, for an oblate spheroid $a$ and $c$ correspond to the semi-major and semi-minor axis, respectively; for an illustration see Supplementary Figure 4A.

Figure 4B shows the perimeter ratio for prolate (blue) and oblate (red) spheroids as a function of the semi-major axis ($c$ for prolate and $a$ for oblate). Due to spherical symmetry, the perimeter ratio between long- and short-axis polarisation is equal to 1 for a sphere. For small deviations from spherical geometry (semi-major axis comparable with the radius of the sphere $R = 18.25\mu m$), the perimeter ratios in the prolate and oblate geometries are nearly the same. For larger deviations, however, the perimeter ratio for a prolate geometry becomes significantly smaller than for an oblate geometry. This difference suggests that long-axis polarisation is more favourable for a prolate spheroid than for an oblate spheroid.

*b.* ***Axis selection for an oblate spheroid.*** As a representative example we analyzed pattern formation in an oblate spheroid with axes $35\,\mu m - 35\,\mu m - 13.2\,\mu m$ and the same volume as a sphere with radius $R = 18.25\,\mu m$ corresponding to a perimeter ratio of 0.72; note the smaller perimeter ratio 0.52 for a prolate spheroid with the same volume and semi-major axis $35\,\mu m$. We performed an extensive set of FEM simulations sweeping both $\lambda$ and $D_{\text{cyt}}$ in a range between $0.01\,s^{-1} - 0.3\,s^{-1}$ (with step size $0.01\,s^{-1}$) and $1.0\,\mu m^2 s^{-1} - 20\,\mu m^2 s^{-1}$ (with step size $1\,\mu m^2 s^{-1}$), respectively, and determined the steady state solution of the reaction-diffusion model. Figure 5A shows a "phase diagram" indicating the parameter regimes where the polarisation axis is oriented along the long or short axis or along some intermediate axis (diagonal). We find that there is indeed a parameter regime where short-axis polarisation is stable, namely for $D_{\text{cyt}}$ smaller than approximately $5\,\mu m^2 s^{-1}$ and independent of the value of $\lambda$. This suggests that weak cytosolic flows are required for stable short-axis polarisation. Interestingly, there is no direct transition between stable short-axis and stable long-axis polarisation but an intermediary regime where the stable polarisation axis is aligned at an intermediate orientation. This indicates a subtle interplay between interface length minimisation and effects due to bulk-to-boundary ratios in this region of the $\lambda - D_{\text{cyt}}$ parameter space.

*c.* ***Axis selection for a prolate spheroid.*** We have just learned that for a large perimeter ratio in an oblate spheroid one can find parameter regimes where short-axis polarisation is stable. However, for the prolate spheroid with the same volume (axes $15\mu m - 15\mu m - 27\mu m$) we only find metastable short-axis polarisation (see section *three-dimensional cell geometry and the role of interface length* and Fig. 6 in the main text). We hypothesize that this is due to the smaller perimeter ratio if compared to an oblate with the same volume (see Supplementary Figure 4).

It is, however, not clear whether short-axis polarisation is always metastable in any prolate spheroids. If the perimeter ratio is indeed an important factor, it should be possible to find

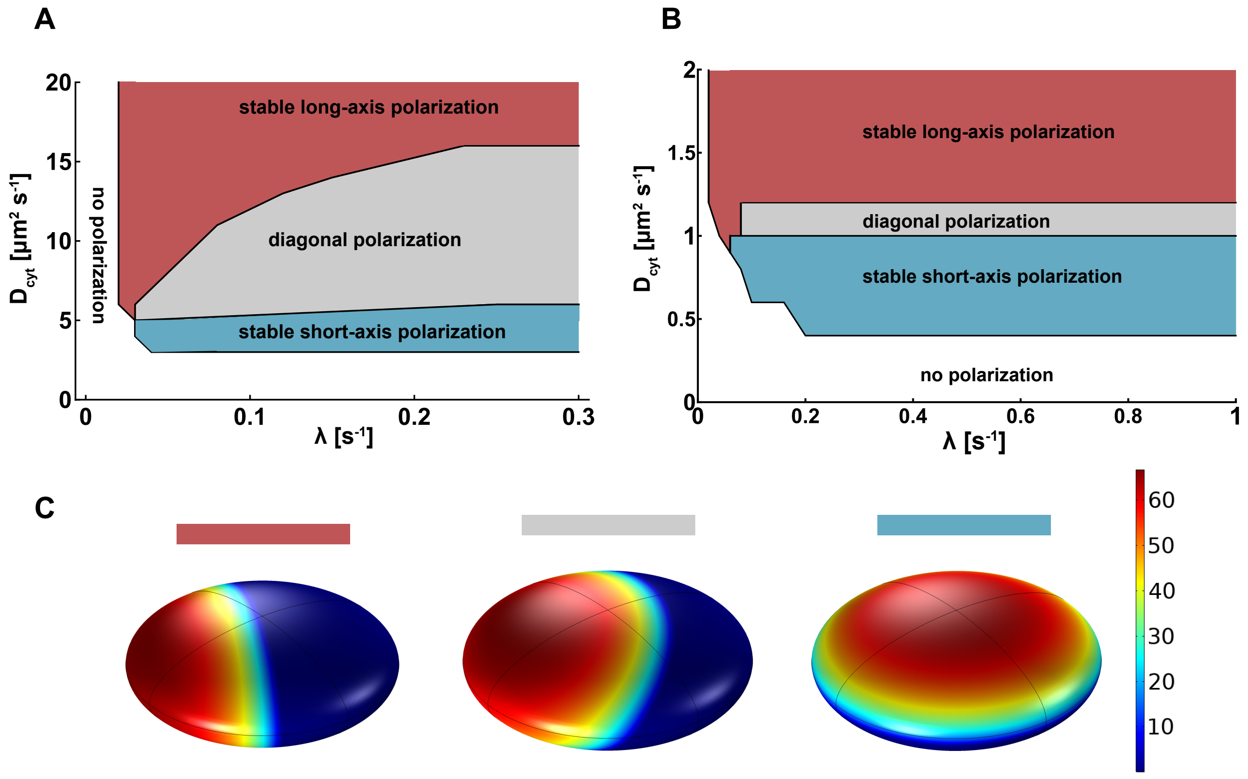


Supplementary Figure 5. Axis selection for oblate and prolate spheroids. Stable polarisation axis in steady state as obtained from FEM simulations for an oblate (**A**) and a prolate spheroid (**B**) in the $\lambda - D_{\text{cyt}}$ parameter space. For an oblate spheroid (**A**), we find that short-axis polarisation is stable for small values of $D_{\text{cyt}}$ (shaded cyan region) quite independent of the value for $\lambda$, while long-axis polarisation is stable for sufficiently large $D_{\text{cyt}}$ and small $\lambda$ (shaded red region), similar to our findings for two-dimensional ellipses (Fig. 3 in the main text). The transition from stable short-axis polarisation to long-axis polarisation is not abrupt but there is an intermediary region where the pattern aligns along the diagonal (shaded grey region). For a prolate spheroid (**B**), we find similar results but for different parameter regimes. Long-axis polarisation is stable for sufficiently large $D_{\text{cyt}}$ (shaded red region), and the regime with diagonal polarisation is less pronounced. (**C**) Typical steady state patterns as obtained from the corresponding parameter combinations in **(A)** and **(B)** (red, grey, and cyan shaded area) shown for an oblate spheroid.

stable short-axis polarisation for a prolate spheroid that has a perimeter ratio comparable with an oblate spheroid (as is the case for prolate spheroids that are almost spherical, cf. Supplementary Figure 4B). To test this, we performed an extensive set of FEM simulations for a prolate spheroid with axes $16.6\,\mu m - 16.6\,\mu m - 22\,\mu m$, corresponding to a perimeter ratio of 0.86; note that an oblate with the same volume and semi-major axis $22\,\mu m$ gives a perimeter ratio of 0.88. We used parameter for $\lambda$ and $D_{\text{cyt}}$ ranging between $0.01s^{-1} - 1.0s^{-1}$ (with step size $0.01s^{-1}$) and $0.4\mu m^2 s^{-1} - 3\mu m^2 s^{-1}$ (with step size $0.2\mu m^2 s^{-1}$). Similar to the oblate case we indeed find that short-axis polarisation can be stabilized for a small parameter region in the $\lambda - D_{\text{cyt}}$ space and that the two regions (stable long- and short- -axis polarisation) are connected by a regime where the pattern aligns along the diagonal (Supplementary Figure 5B). The parameter range for such an intermediate polarisation is, however, significantly smaller as for the oblate case.

### Supplementary Note 6

**Minimisation of the average net cytosolic protein flux onto the membrane explains interface minimisation.** To shed more light on the observed interface minimisation in three-dimensional ellipsoidal geometries we analysed the net cytosolic protein fluxes onto the membrane for the different pPAR and aPAR protein species:

$$J_{\text{net}}^{(\text{P})} = D_{\text{cyt}}\nabla_{\perp}c_P + D_{\text{cyt}}\nabla_{\perp}\, c_{P^*} = k_p^{\text{off}}\, m_P - k_p^{\text{on}}\, c_P + k_{Pa}\, m_P\, m_{A_{12}}\,, \tag{1}$$

$$\begin{aligned} J_{\text{net}}^{(\text{A}_1)} &= D_{\text{cyt}}\nabla_{\perp}c_{A_1} + D_{\text{cyt}}\nabla_{\perp}c_{A_1^*} \\ &= k_a^{\text{off}}\left(m_{A_1} + m_{A_{12}}\right) - k_a^{\text{on}}c_{A_1} + k_{Ap}\, m_P\left(m_{A_1} + m_{A_{12}}\right)\,, \end{aligned} \tag{2}$$

$$J_{\text{net}}^{\text{A}_2} = D_{\text{cyt}}\nabla_{\perp}c_{A_2} = \left(k_{Ap}\, m_P + k_a^{\text{off}}\right) m_{A_{12}} - k_d c_{A_2} m_{A_1}\,, \tag{3}$$

see also in the Methods Section the paragraph on reactive boundary conditions.

Strikingly, we find that all of the local net protein fluxes $J_{\text{net}}^{(\text{P}/\text{A}_{1,2})}$ remain constant as the pattern rotates from short- to long-axis polarisation; as an example the pPAR flux is shown in Supplementary Figure 6. Hence, one expects that the averages of the absolute values of the net membrane fluxes integrated over the whole membrane area $\partial\Omega$

$$\overline{J}^{(\text{P}/\text{A}_{1,2})} = \int_{\partial\Omega} |J_{\text{net}}^{(\text{P}/\text{A}_{1,2})}|\, \text{d}S \Big/ \int_{\partial\Omega} \text{d}S \tag{4}$$

are expected to be larger for short-axis polarisation than for long-axis polarisation, simply due to the larger interface perimeter. This is indeed the case: for the pPAR flux shown in Supplementary Figure 6, we find that the average absolute net flux ratio between long- and short-axis polarisation is $\overline{J}_{\text{long}}^{(\text{P})}/\overline{J}_{\text{short}}^{(\text{P})} = 0.66$. This indicates that long-axis polarisation is maintained by a smaller total protein flux and is therefore more favourable.

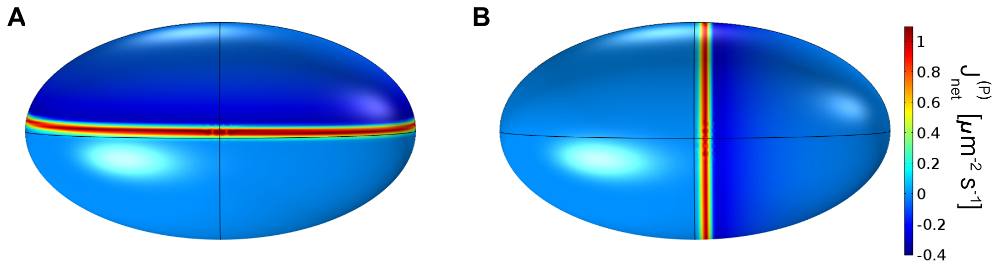


Supplementary Figure 6. Illustration of protein fluxes onto the membrane. (**A**) A snapshot of the net flux $J_{\text{net}}^{(\text{P})}$ of pPAR proteins where the system is polarised in a metastable state (short-axis polarisation) is shown. (**B**) A snapshot of the net flux $J_{\text{net}}^{(\text{P})}$ of pPAR proteins where the system is polarised in a long-axis polarised is shown. The net flux of pPAR proteins along the interface has the same local magnitude for the steady state with long-axis polarisation as for the metastable short-axis polarisation. All parameters are set as in Table 6.

### Supplementary Note 7

**Cytosolic fluxes depend on the cytosolic diffusion and dictate the transition time from short to long axis polarisation.** As discussed in the main text and shown there in Fig. 6, the transition time from short- to long-axis polarisation (for a 3d prolate spheroid) depends on both the reactivation rate $\lambda$ and the cytosolic diffusion $D_{\text{cyt}}$. However, this dependence is not simply explained by the reactivation length $\ell$ alone, since our results show that actually the dependence on the cytosolic diffusion constant $D_{\text{cyt}}$ is decisively stronger than that on $\lambda$. Because the transition from short- to long-axis polarisation (interface minimisation) is driven by protein fluxes, we investigated the cytosolic protein flux for different cytosolic diffusion constants $D_{\text{cyt}}$.

Figure 7A shows the magnitude of the cytosolic flux of species $A_1$ after the steady state (long-axis polarisation) has been reached. We defined the magnitude of the cytosolic flux as its Euclidean norm:

$$||\mathbf{J}_{A_1}|| = D_{\text{cyt}}\, ||\big(\partial_x c_{A_1}, \partial_y c_{A_1}, \partial_z c_{A_1}\big)||\,. \tag{5}$$

This flux decreases with increasing distance from the membrane. Moreover, the lower the

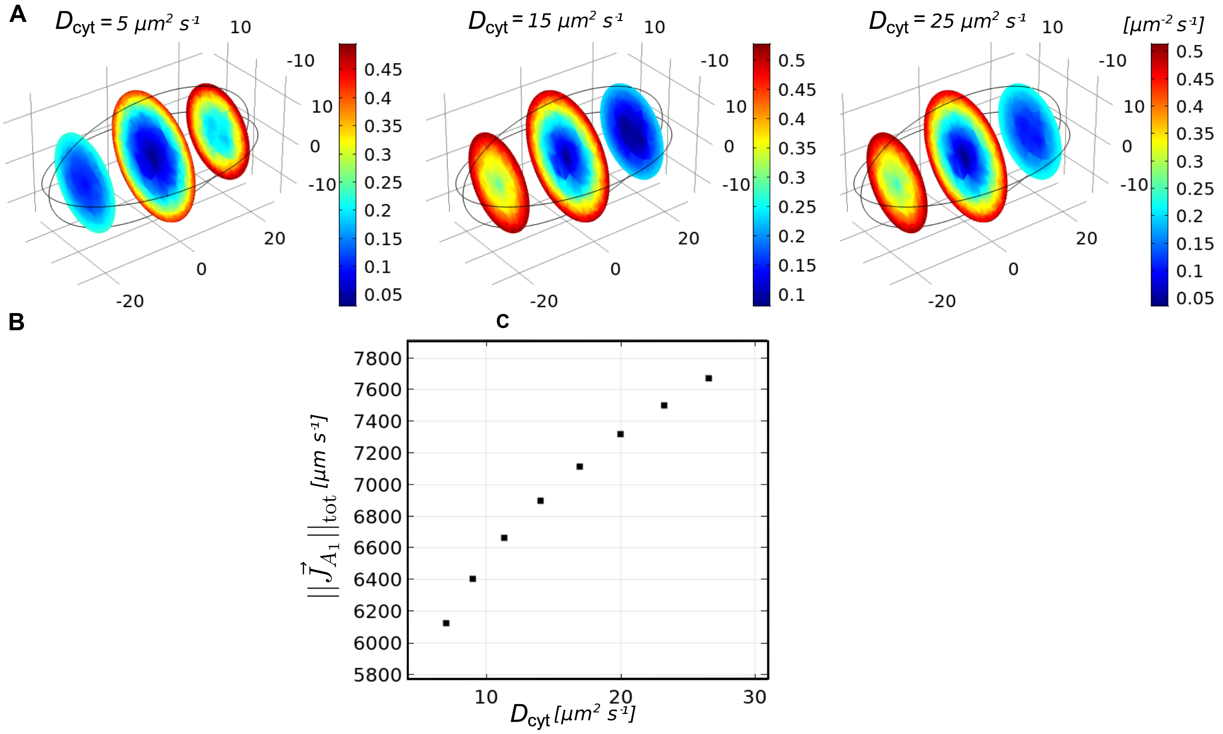


Supplementary Figure 7. Illustration of cytosolic fluxes. (**A**) The magnitudes of cytosolic fluxes of species $A_1$ for three different cytosolic diffusion constants (indicated in the graph) are shown for three slices through the cytosol at $x = 0\mu m$ and $x = \pm 18\,\mu m$. The reactivation rate was set to $\lambda = 0.15 s^{-1}$ and all other parameters were set as given in the main text table 4. (**B**) The overall cytosolic flux (absolute value of flux integrated over the full cytosolic volume) is shown as a function of the cytosolic diffusion constant.

cytosolic diffusion the steeper are the flux gradients, i.e. the shorter is the penetration depth of the flux from the membrane into the cytosol; the width of the red domains (at midcell) in Supplementary Figure 7A decreases with lowering the diffusion constant from $D_{\text{cyt}} = 25\,\mu m^2 s^{-1}$ to $5\,\mu m^2 s^{-1}$. We also notice that the polar cytosolic region shows high cytosolic fluxes on the pPAR-side of the cell, i.e. where the $P$ domain is on the membrane. In contrast, the cytosolic flux of $A_1$ is very low (blue in Supplementary Figure 7A) in the polar region where $A_1$ builds the domain on the membrane. Figure 7B shows the magnitude of the cytosolic flux of species $A_1$ integrated over the whole cytosol (total flux)

$$||\mathbf{J}_{A_1}||_{\text{tot}} = D_{\text{cyt}} \int_{\Omega} ||(\partial_x c_{A_1}, \partial_y c_{A_1}, \partial_z c_{A_1})|| \tag{6}$$

as a function of the cytosolic diffusion constant. Clearly, with increasing cytosolic diffusion constant, the overall cytosolic flux is increasing. Together with the observation that the transition times become shorter with increasing cytosolic diffusion constant (see Fig. 6D in the main text) this shows that there is a correlation between faster transition times and higher cytosolic fluxes.

### Supplementary Note 8

**Time scales for the formation of cell polarisation.** In order to determine the time required for the formation of long-axis polarisation, we consider an idealised situation where this is achieved by the PAR reaction-diffusion system alone. For the cell polarisation process in *C. elegans* there is experimental evidence that the localisation of the centrosome as well as the successive actomyosin contraction play an important role in polarity establishment and support its alignment with the long axis ( [4, 6, 7]). However, how the PAR reaction-diffusion system acts in concert with actomyosin contraction is not understood in realistic three--dimensional cell geometry. Previous work uses a simplified one-dimensional cell geometry ( [1, 4, 8]). Here, we focus (as a first and important step) on the reaction-diffusion pathway alone disregarding any effects due to the PAR interaction with the centrosome or actomyosin contraction and ensuing cytoplasmic flows. This way one can learn how robust and fast reaction-diffusion dynamics on its own can establish long-axis polarisation and what the relative role of other effects like cytoplasmic flow may be. In the actual C. elegans embryo polarisation has to be stable along the long axis for $\approx 15\,\text{min}$ until the first cell division. Therefore, the time of a possibly existing short- axis polarisation and the transition to the long axis is an important observable for the real system. Hence we ask: How fast is the long axis selected as the stable polarisation axis driven solely by a reaction-diffusion dynamics?

We find that the time scales for the selection and maintenance of different polarisation axes depend on the reactivation rate $\lambda$ and the cytosolic diffusion $D_{\text{cyt}}$ both in a two-dimensional elliptical geometry and in an three-dimensional ellipsoidal geometry (see Fig. 6 in the main text and Supplementary Figure 8). Strikingly, the transition time from short to long axis polarisation is extremely slow in 2d compared to 3d ($\approx 1000\,\text{min}$ in 2d compared to $\approx 100\,\text{min}$ for 3d data: compare Supplementary Figure 8 with Fig.6 in the main text). The transition time from any transient polarisation pattern to a steady state long-axis polarisation pattern may be taken as a proxy for the expected typical time scales of polarisation re-alignment in case of an initially non-aligned cue (e.g. this happens if the centrosome does not localise at the poles initially). Hence, we conclude that for physiological parameters in 2d these times are far too long: a wrong alignment induced by cues or flows can not be corrected by a mechanism based on reaction and diffusion alone. In contrast, our simulations in 3d show that these transition times are short in a broad region of parameter space; compare Supplementary Figure 8 with Fig. 6 in the main text. Therefore, we conclude, that all geometry-sensitive mechanisms of the reaction-diffusion system, as well as the activation--deactivation cycle and interface minimisation, play an important role for cell polarisation in *C. elegans*. Antagonism (of aPARs and pPARs) and recruitment (among aPARs) enables polarisation, fast cytosolic diffusion and the activation-deactivation cycle enable the cell to polarise along the long axis from the beginning on, and interface minimisation always leads to long axis polarisation in the long term. Furthermore, cytosolic diffusion, as it determines the magnitude of fluxes, decisively influences perfect polarity establishment along the long axis on a biologically reasonable time scale. E.g. if the centrosome was originally localised close to mid-cell and would induce an initial polarity alignment with the long axis, fast cytosolic diffusion would rescue such an embryo and polarisation would align with the long axis before cell division.

In contrast to the transition time from any initial polarisation to well aligned long-axis polarisation, the establishment time of the initial polarisation from a homogenous aPAR dominated state on the membrane is strongly dependent on the type of initial perturbations. Take Supplementary Figure1 middle and bottom row as an example in 2d, where polarisation is quickly established with initial gradients (despite of misalignment). The establishment time of any polarisation from a homogenous aPAR dominated state with only a small random initial perturbation is of the order of 30 minutes in 3d and is approximately three times slower in 2d. Therefore, with only a small random initial perturbation the reaction-diffusion system alone does still lead to stable polarisation, but on a time scale that is too slow for the real embryo (see Fig. 8 A for 2d and 3d times).

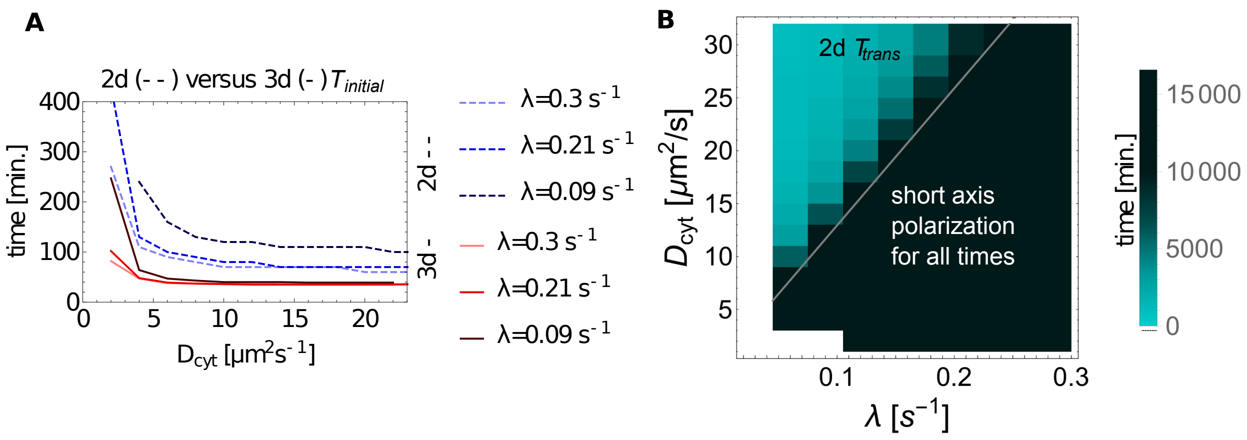


Supplementary Figure 8. **Times in 2d versus 3d**. (**A**) The initial time of polarisation $T_{\text{initial}}$ is plotted against the cytosolic diffusion for various reactivation rates in 2d as well as in 3d. (**B**) $T_{\text{trans}}$ is shown in cyan color code in the $D_{\text{cyt}}$-$\lambda$ parameter space. The gray line shows the line of constant reactivation length, which divides steady state long- and short-axis polarisation $\ell^{\star}$. It was interpolated as a linear function with zero offset.

### SUPPLEMENTARY TABLES

| $k_{Ap}$ | $k_{Pa}$ | steady state | onset |
|---|---|---|---|
| 0.24 | 2.28 | no pattern | transient long axis pol. |
| 0.28 | 2.16 | long axis polarisation | long axis |
| 0.32 | 2.04 | long axis polarisation | long axis |
| 0.36 | 1.92 | long axis polarisation | long axis |
| 0.4 | 1.8 | long axis polarisation | long axis |
| 0.44 | 1.1.68 | long axis polarisation | long axis |
| 0.48 | 1.56 | long axis polarisation | long axis |
| 0.5 | 1.5 | long axis polarisation | long axis |
| 0.52 | 1.44 | long axis polarisation | long axis |
| 0.54 | 1.38 | long axis polarisation | long axis |
| 0.56 | 1.32 | long axis polarisation | long axis |
| 0.58 | 1.26 | long axis polarisation | long axis |
| 0.6 | 1.2 | long axis polarisation | long axis |
| 0.64 | 1.08 | long axis polarisation | long axis |
| 0.68 | 0.96 | long axis polarisation | long axis |
| 0.72 | 0.84 | no pattern | transient long axis pol. |
| 0.76 | 0.72 | no pattern | transient long axis pol. |

Supplementary Table 1. **Sweep of antagonistic rates to investigate the excitable region with initial gradients for slow reactivation.** FEM sample sweeps of $k_{Ap}$, $k_{Pa}$ with initial linear gradient for $\lambda = 0.05s^{-1}$ (for more details see supplementary section A). The sweep shows that also outside of the spontaneously polarizing region the system can be excited into stable long axis polarisation. All other parameters were set as in the standard parameter set shown in Table 1 of the main text.

| $k_{Ap}$ | $k_{Pa}$ | steady state | onset |
|---|---|---|---|
| 0.2 | 2.4 | no pattern | transient short axis p. |
| 0.24 | 2.28 | short axis polarisation | short axis |
| 0.28 | 2.16 | short axis polarisation | short axis |
| 0.32 | 2.04 | short axis polarisation | short axis |
| 0.36 | 1.92 | short axis polarisation | short axis |
| 0.4 | 1.8 | short axis polarisation | short axis |
| 0.44 | 1.1.68 | short axis polarisation | short axis |
| 0.48 | 1.56 | short axis polarisation | short axis |
| 0.5 | 1.5 | short axis polarisation | short axis |
| 0.52 | 1.44 | short axis polarisation | short axis |
| 0.54 | 1.38 | short axis polarisation | short axis |
| 0.56 | 1.32 | short axis polarisation | short axis |
| 0.58 | 1.26 | short axis polarisation | short axis |
| 0.6 | 1.2 | short axis polarisation | short axis |
| 0.64 | 1.08 | short axis polarisation | short axis |
| 0.68 | 0.96 | short axis polarisation | short axis |
| 0.72 | 0.84 | no pattern | transient short axis p. |
| 0.76 | 0.72 | no pattern | transient short axis p. |

Supplementary Table 2. **Sweep of antagonistic rates to investigate the excitable region with initial gradients for fast reactivation.** FEM sample sweeps of $k_{Ap}$, $k_{Pa}$ with initial linear gradient for $\lambda = 0.3s^{-1}$ (for more details see supplementary section A). The sweep shows that also outside of the spontaneously polarizing region the system can be excited into stable short axis polarisation. All other parameters were set as in the standard parameter set shown in Table 1 of the main text.

| $k_{Ap}$ | $k_{Pa}$ | steady state | onset |
|---|---|---|---|
| 0.2 | 2.4 | no pattern | transient short axis p. |
| 0.24 | 2.28 | short axis polarisation | short axis |
| 0.28 | 2.16 | short axis polarisation | short axis |
| 0.32 | 2.04 | short axis polarisation | short axis |
| 0.36 | 1.92 | short axis polarisation | short axis |
| 0.4 | 1.8 | short axis polarisation | short axis |
| 0.44 | 1.1.68 | short axis polarisation | short axis |
| 0.48 | 1.56 | short axis polarisation | short axis |
| 0.5 | 1.5 | short axis polarisation | short axis |
| 0.52 | 1.44 | short axis polarisation | short axis |
| 0.54 | 1.38 | short axis polarisation | short axis |
| 0.56 | 1.32 | short axis polarisation | short axis |
| 0.58 | 1.26 | short axis polarisation | short axis |
| 0.6 | 1.2 | short axis polarisation | short axis |
| 0.64 | 1.08 | short axis polarisation | short axis |
| 0.68 | 0.96 | short axis polarisation | short axis |
| 0.72 | 0.84 | no pattern | transient short axis p. |
| 0.76 | 0.72 | no pattern | transient short axis p. |

Supplementary Table 3. **Sweep of antagonistic rates to investigate the excitable region with initial gradients for fast reactivation.** FEM sample sweeps of $k_{Ap}$, $k_{Pa}$ with initial linear gradient for $\lambda = 1.s^{-1}$ (for more details see supplementary section A). The sweep shows that also outside of the spontaneously polarizing region the system can be excited into stable short axis polarisation. All other parameters were set as in the standard parameter set shown in Table 1 of the main text.

| Parameter | Value |
|---|---|
| $a$ | $13.2\,\mu m$ |
| $b$ | $13.2\,\mu m$ |
| $c$ | $35\,\mu m$ |
| $k_{a/p}^{on}$ | $0.1\,\mu m \cdot s^{-1}$ |
| $k_{a/p}^{off}$ | $0.005\,s^{-1}$ |
| $k_{Ap}$ | $0.4\,\mu m^2 \cdot s^{-1}$ |
| $k_{Pa}$ | $1.2\,\mu m^2 \cdot s^{-1}$ |
| $k_d$ | $0.15\,\mu m^3 \cdot s^{-1}$ |
| $D_{mem}^a$ | $0.28\,\mu m^2 \cdot s^{-1}$ |
| $D_{mem}^p$ | $0.15\,\mu m^2 \cdot s^{-1}$ |
| $\rho_{A1}$ | $10.5\,\mu m^{-3}$ |
| $\rho_{A2}$ | $2.5\,\mu m^{-3}$ |
| $\rho_P$ | $12.0\,\mu m^{-3}$ |

Supplementary Table 4. Parameter set for the oblate 3d FEM sweep in Fig. 5. All parameters were fixed to the values shown above except for $D_{\text{cyt}}$ and $\lambda$. The cytosolic difffusion constant $D_{\text{cyt}}$ was varied between $1.0\,\mu m^2 \cdot s^{-1} - 20\,\mu m^2 \cdot s^{-1}$ (with step size of $1\,\mu m^2 \cdot s^{-1}$) and the reactivation rate $\lambda$ was varied between $0.01\,s^{-1} - 0.3\,s^{-1}$ (with step size of $0.01\,s^{-1}$) to generate the result shown in Fig 5.

| Parameter | Value |
|---|---|
| $a$ | $22\,\mu m$ |
| $b$ | $22\,\mu m$ |
| $c$ | $16.6\,\mu m$ |
| $k_{a/p}^{on}$ | $0.1\,\mu m \cdot s^{-1}$ |
| $k_{a/p}^{off}$ | $0.005\,s^{-1}$ |
| $k_{Ap}$ | $0.4\,\mu m^2 \cdot s^{-1}$ |
| $k_{Pa}$ | $1.2\,\mu m^2 \cdot s^{-1}$ |
| $k_d$ | $0.15\,\mu m^3 \cdot s^{-1}$ |
| $D_{mem}^a$ | $0.28\,\mu m^2 \cdot s^{-1}$ |
| $D_{mem}^p$ | $0.15\,\mu m^2 \cdot s^{-1}$ |
| $\rho_{A1}$ | $10.5\,\mu m^{-3}$ |
| $\rho_{A2}$ | $2.5\,\mu m^{-3}$ |
| $\rho_P$ | $18.0\,\mu m^{-3}$ |

Supplementary Table 5. Parameter set for the prolate 3d FEM sweep in Fig. 5. All parameters were fixed to the values shown above except for $D_{\text{cyt}}$ and $\lambda$. The cytosolic difffusion constant $D_{\text{cyt}}$ was varied between $0.4\,\mu m^2 \cdot s^{-1} - 3.0\,\mu m^2 \cdot s^{-1}$ (with step size of $0.2\,\mu m^2 \cdot s^{-1}$) and the reactivation rate $\lambda$ was varied between $0.01\,s^{-1} - 1.0\,s^{-1}$ (with step size of $0.01\,s^{-1}$) to generate the result shown in Fig 5.

| Parameter | Value |
|---|---|
| $a$ | $27\,\mu m$ |
| $b$ | $15\,\mu m$ |
| $c$ | $15\,\mu m$ |
| $D_{\text{cyt}}$ | $10\,\mu m^2 s^{-1}$ |
| $\lambda$ | $0.2\,s^{-1}$ |
| $k_{a/p}^{on}$ | $0.1\,\mu m \cdot s^{-1}$ |
| $k_{a/p}^{off}$ | $0.005\,s^{-1}$ |
| $k_{Ap}$ | $0.4\,\mu m^2 \cdot s^{-1}$ |
| $k_{Pa}$ | $1.2\,\mu m^2 \cdot s^{-1}$ |
| $k_d$ | $0.034\,\mu m^3 \cdot s^{-1}$ |
| $D_{mem}^{a}$ | $0.28\,\mu m^2 \cdot s^{-1}$ |
| $D_{mem}^{p}$ | $0.15\,\mu m^2 \cdot s^{-1}$ |
| $\rho_{A1}$ | $10.5\,\mu m^{-3}$ |
| $\rho_{A2}$ | $2.5\,\mu m^{-3}$ |
| $\rho_{P}$ | $8.0\,\mu m^{-3}$ |

Supplementary Table 6. Parameter set for the pPAR average net membrane flux shown in Fig. 6.

### SUPPLEMENTARY REFERENCES

[1] Goehring, N. W. *et al.* Polarization of PAR proteins by advective triggering of a pattern-forming system. *Science* **334**, 1137–1141 (2011).

[2] Goehring, N. W., Hoege, C., Grill, S. W. & Hyman, A. PAR proteins diffuse freely across the anterior-posterior boundary in polarized C. elegans embryos. *J Cell Biol* **193**, 583–594 (2011).

[3] Robin, F. B., McFadden, W. M., Yao, B. & Munro, E. M. Single-molecule analysis of cell surface dynamics in Caenorhabditis elegans embryos. *Nat Methods* **11**, 677–682 (2014).

[4] Gross, P. *et al.* Guiding self-organized pattern formation in cell polarity establishment. *Nat Phys* **15**, 293–300 (2019).

[5] Halatek, J. & Frey, E. Rethinking pattern formation in reaction-diffusion systems. *Nat Phys* **14**, 507–514 (2018).

[6] Nance, J. & Zallen, J. Elaborating polarity: PAR proteins and the cytoskeleton. *Development* 799–809 (2011).

[7] Motegi, F. & Seydoux, G. The PAR network: redundancy and robustness in a symmetry-breaking system. *Philos Trans R Soc Lond B Biol Sci* **368**, 20130010 (2013).

[8] Dawes, A. T. & Munro, E. M. PAR-3 oligomerization may provide an actin-independent mechanism to maintain distinct par protein domains in the early Caenorhabditis elegans embryo. *Biophys J* **101**, 1412–1422 (2011).
